# Environmental influences and individual characteristics that affect learner-centered teaching practices

**DOI:** 10.1101/2020.09.21.306407

**Authors:** Nathan Emery, Jessica Middlemis Maher, Diane Ebert-May

## Abstract

Research-based teaching practices can improve student learning outcomes in a variety of complex educational environments. The implementation of learner-centered teaching practices in STEM can both benefit or be constrained by different factors related to individual instructors and the teaching environment. However, we know little of how the instructional climate varies across institutions and how this climate affects teaching practices. Our study sought to describe the relative importance of environmental influences and individual characteristics on learner-centered teaching practices across institutions. We also assessed departmental climate for 35 US higher education institutions. We found that self-efficacy in teaching and professional development exert a strong influence on faculty teaching practices. While departmental climate did not emerge as a significant predictor of teaching practices, there was consistently low support for teaching, and institution size was negatively correlated with leadership and evaluation of effective teaching. We also found that professional development may prepare instructors to teach learner-centered courses in different collegial teaching climates. Our results suggest that through cultivating self-efficacy and participating in iterative professional development, instructors can implement effective teaching practices across institutional environments.

## Introduction

As scientific research and knowledge continues to grow, the need to provide undergraduate students the opportunity to learn science is more urgent than ever [1, 2]. A convincing body of evidence indicates that teaching strategies to promote active student engagement with science are most likely to increase student mastery of concepts and scientific practices and reduce achievement gaps in higher education [3–5]. Research-based pedagogical strategies can positively influence student learning outcomes as compared to traditional lecture-based approaches in which there is minimal student interaction with the instructor or each other. Changes in how science is taught also reduces achievement gaps and promotes retention in the discipline [6, 7].

In complex higher education systems, there are many factors that may affect the implementation of learner-centered teaching practices in STEM courses. Instructors come from a range of pedagogical backgrounds [8], confidence levels [9], and personal beliefs and intentions about teaching [10]. Teaching practices are also influenced by personal experiences [11], peer interactions [12], and professional development [13]. In higher education, instructors are faced with overlapping professional obligations and teaching support and constraints from their institution, department, and peers [14, 15]. While past research has explored some of these influences on teaching practices, few studies have taken a multidimensional approach to exploring the factors that influence teaching of learner-centered courses [16–18]. As part of a longitudinal study of the Faculty Institutes for Reforming Science Teaching (FIRST) IV program [19, 20], we sought to tease apart the relative factors that drive the implementation of learner-centered teaching practices. In [20] we found that teaching professional development outcomes persisted in the long-term and across a career transition (from postdoc to faculty). However, we still do not know why individual instructors are teaching more or less learner-centered courses at their respective institutions. Assessment of teaching professional development programs allows us to identify the variables that are most closely associated with changes in teaching, specifically the potential factors that facilitate or constrain learner-centered teaching practice. Previous studies of teaching professional development have argued for a variety of interacting factors playing a role in teaching practices [9,13,21–23]. Social cognitive theory posits that personal factors and environmental influences interact with and determine behavior [24].

Inspired by this theory, we sought to uncover the relative impact of individual characteristics and environmental influences on learner-centered teaching practices in higher education.

### Individual Characteristics

As an instructor, there are a variety of characteristics, experiences, and beliefs that could affect teaching practices in the classroom. While knowledge of and experience in learner-centered practices can translate into the classroom [25], individuals are also influenced by their current beliefs and intentions surrounding teaching [10]. It is well established that beliefs and attitudes can affect behavior [26]. Teacher-focused beliefs and approaches can hinder the adoption and implementation of learner-centered teaching practices [27]. On the other hand, approaching courses with the intent to facilitate learning among students can manifest in learner-centered teaching practices [20].

Self-efficacy, the confidence in one’s ability to perform a task, could also play a role in teacher practices in the classroom. Perceived self-efficacy is a universal construct [28, 29] and is particularly important to consider for faculty participating in teaching and research [30–32]. High teaching self-efficacy is particularly important for instructors in higher education classrooms [9,33,34]. While there are likely many ways that faculty gain self-efficacy [35, 36], there is some evidence that professional development can lead to increases in teaching self-efficacy [33,37,38]. Not only does high teaching self-efficacy benefit the instructor, but it also could have positive impacts on students and student learning outcomes. Given the consistent effect of self-efficacy on task performance in the literature [31, 39], it is likely that teaching self-efficacy affects teaching practices in the classroom.

### Environmental Influences

While there are many characteristics that define how an individual instructor teaches, their teaching practice has likely been influenced by environmental factors throughout their career. These include, but are not limited to, faculty time allocation, course characteristics, professional development experiences, and the organizational structure and climate surrounding teaching. Faculty responsibilities can vary. The extent to which a faculty member dedicates their time to research, teaching, and service depends on departmental guidelines, tenure and promotion criteria, and personal choice [40]. Time allocation for teaching may differ across faculty and be reflected in their teaching practices. Past research has shown evidence of teaching differences in contingent/part-time faculty and tenure-track/tenured faculty [41–43]. Teaching focused faculty, while hired primarily for their teaching role, end up participating in research and service and serve similar roles at institutions as their research-focused colleagues [44, 45]. While student learning outcomes appear to be similar between tenure-track teaching faculty and tenure-track research faculty, teaching faculty carry a larger teaching requirement than other faculty [46]. Because of the pressures on faculty to spend more time on teaching or research [40, 47], faculty time allocation could affect teaching practices in the classroom.

Another possible constraint to learner-centered teaching is the course itself. Specific course types or sizes can facilitate particular teaching approaches or practices. Large enrollment courses tend to operate under lecture format [4, 48], with the instructor as the sole source of information and knowledge. Large courses also reduce the frequency and quality of feedback and interaction between faculty and students [3,49,50]. However, recommendations and guidance have been published on how to engage students in large classrooms [51–54]. While course size may incorporate multiple components of course constraints on teaching practices, several other course characteristics have been cited to affect teaching practices including classroom space/design [55, 56], access to technology [57], and student characteristics [58].

Professional development and training can have immediate positive effects on teacher approaches and practices in the classroom [23,33,59,60]) and long-term effects [20,61–63]. By shifting faculty attitudes and approaches to teaching, professional development programs can affect change in the classroom [23,33,64,65]. While these programs work with individuals and have shown to shift faculty teaching practices, it is important to consider the teaching environment. Past literature has advocated that frameworks for evaluating teacher development programs take into account institutional policies and culture surrounding teaching [19, 66].

Organizational structure and climate can vary greatly with respect to teaching and learning in higher education [67, 68]. This variation can lead to differing levels of departmental support, funding and resources, and interest and buy-in from faculty with respect to learner-centered teaching practices. In academic settings, there are multiple levels of organizational structure, from department to institution. Each level has different incentives and pressures for faculty, in particular, departments tend to have significant control over instruction and curriculum design [14, 27]. Departments can both hinder or facilitate implementation of learner-centered practices [69]. They are also the level in which faculty have the most social/professional interactions [70, 71] that may influence their teaching approaches and practices. A positive departmental (professional) culture around teaching can lead to more supportive conversations about teaching and learning [74]. The multiple components of departmental climate with respect to teaching: leadership, support, resources, respect, and faculty-faculty interactions comprise the instructional climate within a department [75]. Several studies have quantified the instructional climate of departments [75, 76], however, much less is known about how climate varies across institutions.

### Research Questions

With little known about the variation in instructional climates and the interacting factors that drive faculty teaching practices, we sought to address the following research questions: 1. What is the relative importance of environmental influences and individual characteristics on learner-centered teaching practices? 2. How does instructional climate vary across institutions?

To answer these questions, we used a linear model and model selection for several variables we hypothesized to affect learner-centered teaching practices (Fig 1). We found that self-efficacy in teaching and professional development were relatively important for faculty implementation of learner-centered teaching in the classroom. We also found that aspects of departmental climate were correlated with institution size and resources and support for

**Figure 1.**
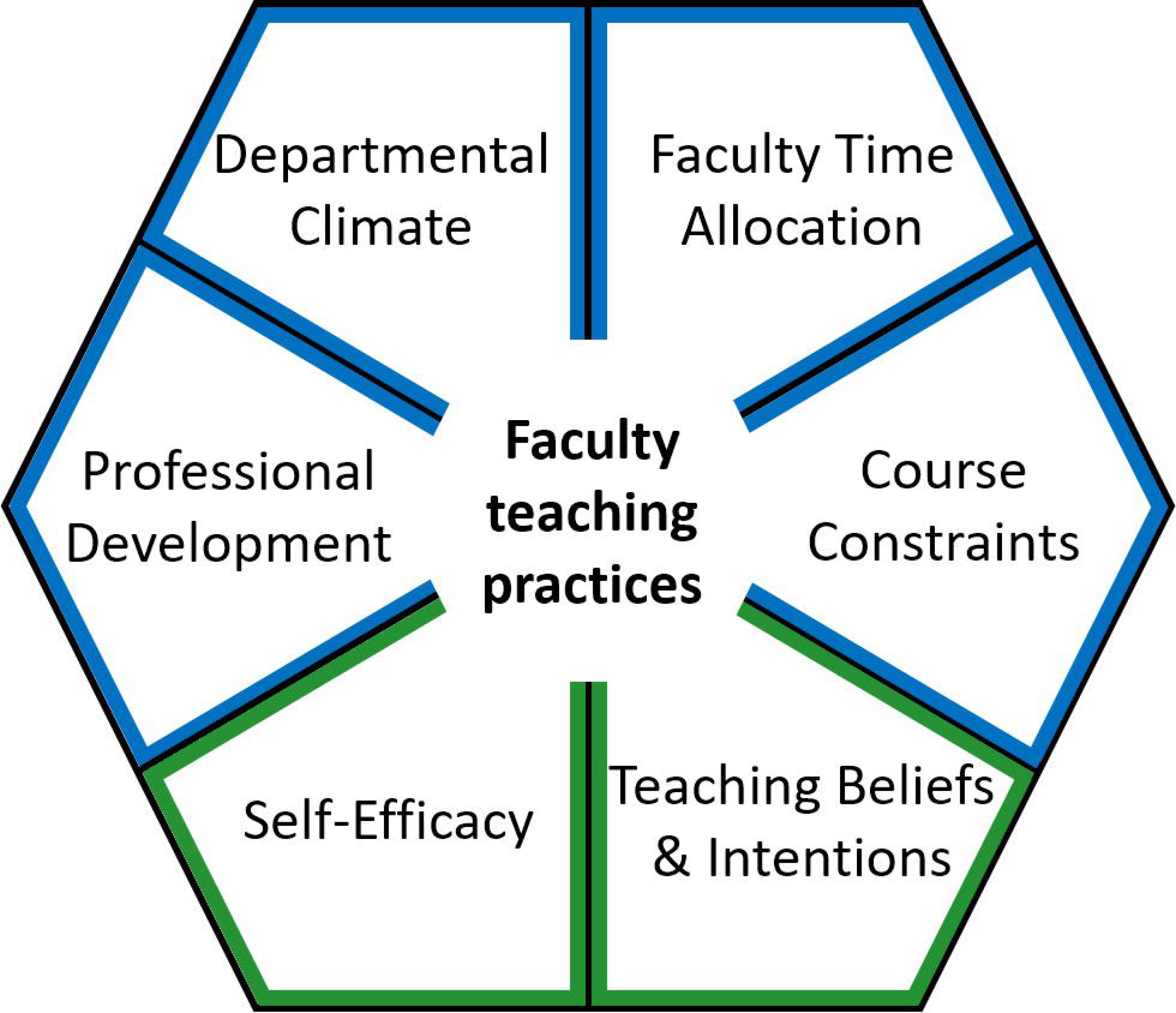
A model of faculty teaching practices as affected by individual characteristics (green) and outside influences (blue). Specific metrics or subscales used in the multivariate model of learner-centered teaching practices included participation in the FIRST IV program (Professional Development), Support for teaching (Departmental Climate), Percent time spent teaching (Faculty Time Allocation), Course size (Course Constraints), Self-efficacy in teaching (Self-Efficacy), and Intentions with regards to knowledge transmission (Teaching Beliefs & Intentions).

## Methods

### Study Design

Our three-year longitudinal study assessed faculty at 35 institutions across the United States [19]. The institutions consisted of doctoral universities, master’s colleges, baccalaureate colleges, and community colleges. Forty faculty were former FIRST IV program participants who then recruited a paired colleague at a similar career stage in the same department. We developed a paired comparison design to assess potential differences in faculty who had experienced significant professional development and their peers. We attempted to reduce the relative influence of institutional/departmental culture by having each pair come from the same department. All participants were given an honorarium for their commitment and contributions to the study.

### Data Collection

At the beginning of the study, each participant completed a survey to provide demographic information and report their knowledge and experience surrounding learner-centered teaching, their perceptions of departmental culture, challenges to implementing active learning, and past teaching experiences (See S1). Then for three years, we assessed faculty teaching approaches [77], teaching practices [78], and student perceptions of the classroom [79] as examined in past work [20]. We also surveyed faculty using instruments on teaching beliefs and intentions [10] and teaching self-efficacy [33].

From 2009-2013, FIRST IV postdoctoral fellow participants (n = 201) selected from institutions nationwide engaged in a two-year teaching professional development program based on learning theory and the principles of evidence-based instructional strategies. As teams, the postdocs developed learner-centered teaching practices, designed an entire course using those principles, and taught the courses during the academic year following the first and second summer workshops at their respective institutions. Specifically, during year one, the postdocs completed a 4-day summer workshop in which they learned to actively engage students in both large and small enrollment biology courses, use individual and group learning strategies, use backward design, and write assessments that were aligned with learning objectives and instruction. Key to the first workshop was establishing teams of postdocs who designed an entire learner-centered, introductory biology course based on how students learn and teaching using scientific practices (for example, creating and testing models, designing arguments, working with data) to help students and do something with core concepts. Following workshop one, the postdocs completed a teaching experience in their home department or in another unit on campus. The teaching experience was mentored by team leaders in the project who served as remote teaching mentors for each group of postdocs. During the second workshop in year two (3 or 4 days) postdocs reflected on and discussed the challenges they encountered during their teaching mentors, identified strengths and elements of instruction that needed work. Postdocs analyzed their course design, which was now informed with student assessment data from their teaching during the previous academic year, and continued to revise their entire course. During the second academic year, the postdocs completed a second teaching experience (a full or partial course), and some taught the course that they had developed in the context of a new faculty position. Again, long-distance teaching mentors and a private FIRSTIV email listserv were valuable tools for maintaining contact and providing immediate support to these early-career instructors. For more details about the program see [13].

All teaching videos recorded by participants from 2009-2013 (historical) and 2016-2019 (current study) were scored with the Reformed Teaching Observation Protocol (RTOP) [78]. As per methods described in [20], two videos per course each year for a total of 6 per participant were collected for the current study, and each video was rated by independent, doctoral-trained biologists from several institutions. Over the duration of the study, 12 raters were involved in viewing videos. The raters were calibrated to one another with an intra-class correlation coefficient of 0.77 over 6 videos. We randomly assigned each de-identified teaching video to two raters and the video scores were averaged per participant for either historical or current results (n=4 minimum per participant; total 400 plus videos). Any video scores with a standard deviation above seven were assigned to a third rater, and the subsequent two closest scores were used in the analyses.

### Teaching Beliefs and Intentions

An important aspect of how faculty teach is their beliefs about teaching practices [80]. To assess faculty beliefs and intentions about teaching, participants completed the Teaching Beliefs and Intentions (TBI) instrument in the third year of the study [10]. This survey instrument consists of 34 items and four subscales: “Beliefs: Learning facilitation”, “Beliefs: Knowledge transmission”, “Intentions: Learning facilitation”, and “Intentions: Knowledge transmission.” The subscales associated with “Knowledge transmission” refer to an instructor’s orientation towards passing information to students directly through instruction/lecturing. “Learning facilitation” is more associated with guiding students through the construction of their own knowledge. The intentions quantified by this instrument are reflective of a teacher’s conceptions about teaching and their academic and social context [10].

### Self-efficacy

How an instructor conducts themselves in class and teaches a course can be partially attributed to self-efficacy, or confidence in their ability [33, 81]. Faculty in our study were assessed on their teaching self-efficacy using a published instrument [33]. This instrument consists of 23 items and six subscales: “Course planning,” “Teaching methods,” “Creating learning environment,” “Assessing student learning,” “Interacting with students,” and “Mastering subject knowledge.” This instrument was distributed to faculty participants in the beginning of their course in the second year of the study.

### Teaching Practice

Observations of teaching provide valuable feedback about how an instructor conducts a course. Participating faculty recorded two class sessions in one course for each year of participation. These recordings were uploaded to a cloud server, de-identified by the authors and distributed to raters (with doctoral degrees in biological sciences) trained in the Reformed Teaching Observation Protocol [78]. For detailed methods, see [20]. We averaged the two ratings per video and if the standard deviation was over 7 (half of a RTOP category), we introduced a third rater and chose the two closest ratings. For statistical analysis, all RTOP scores (up to six per participant) were averaged together to represent an overall RTOP score per participant. Faculty that provided fewer than four videos were removed from the model analysis due to past research recommendations [4]. For former FIRST IV participants we calculated the change in teaching practice by subtracting their mean RTOP score for the current study from their mean RTOP score at the end of the FIRST IV program [13].

### Departmental Climate

Faculty in higher education teach courses within an institution and usually within a department. The discourse and interactions among faculty in a department can be influential and ultimately affect teaching practices [82]. To best characterize the teaching climate of each department, we distributed a survey instrument called the Survey of Climate for Instructional Improvement (SCII) [75]. This instrument consists of 30 items and five subscales: “Leadership,” “Collegiality,” “Resources,” “Respect for teaching,” and “Support for teaching.” We contacted the chairs of every department involved in the study and either sent them a link to distribute to faculty (68% of departments), or distributed the link ourselves to all faculty members in the department according to departmental websites. Departmental data were only counted if greater than 20% of the faculty in the department responded to the survey. This resulted in the removal of three departments out of 35 that participated in the survey. One department chair requested the responses of their faculty, which was not within the scope of the IRB, and thus that department did not participate in the climate survey.

To assess reliability of the SCII instrument (30-item, 5 factors) we performed a confirmatory factor analysis using data collected from all departments (35 departments, N = 352 complete responses) and the lavaan R package [83]. A Royston test determined that the data were not normally distributed, so we used the maximum likelihood estimator for the confirmatory factor analysis. A Kaiser-Meyer-Olkin test for sampling adequacy resulted in a “meritorious” designation (0.93). The prescribed factors, according to thresholds described by Harshman and Stains [84], fell within the Poorer Fit designation for x^2^/df (3.68), “root mean square error of approximation” (RMSEA; 0.087), “standardized root mean square residual” (SRMR; 0.078), “comparative fit index” (CFI; 0.83) and “Tucker-Lewis index” (TLI; 0.813).

Since the published factors of the instrument proved to be unreliable for our sample population, we conducted an exploratory factor analysis of our data. A parallel analysis from the Psych R package [85] suggested six factors as appropriate for our survey data. We then performed a factor analysis and established a cut-off of loadings less than 0.3. This resulted in one item being removed from the original survey: “In my department, instructors with a record of teaching excellence are financially rewarded (e.g., bonuses, raises, or similar).” The resulting fit indices were supportive of the six factor structure with a Best fit designation for SRMR (0.057), Better fit for x^2^/df (2.61), RMSEA (0.068), CFI (0.906), and Poorer fit for TLI (0.894). While the Leadership factor from the original instrument was unchanged, all other factors were renamed according to the resorted items that loaded above 0.3 for each factor. The new factors were called “Mentoring and Material Support” (7 items), “Flexibility in Teaching” (2 items), “Collegiality” (4 items), “Evaluation of Effective Teaching” (5 items), and “Support for Teaching Improvement” (4 items). For a detailed list of items associated with each factor and factor loadings, see S2.

### Statistical Analysis

We evaluated differences in background survey items with paired Wilcoxon signed rank tests. This non-parametric test accounts for the nominal data produced from faculty responses.

Agreement with statements encompassing departmental culture surrounding student-centered teaching were on a scale from one to ten. The degree to which an obstacle makes active learning more challenging was rated on a scale from one to five.

The TBI instrument consists of four independent subscales. We tested for differences between faculty groups for each subscale using a paired Wilcoxon signed rank test. This non-parametric test is appropriate for nominal data and takes into account the paired nature of the study.

We tested for differences between faculty groups with regards to teaching self-efficacy using a paired Wilcoxon signed rank test. This non-parametric test accommodates the nominal data from the self-efficacy instrument and the paired nature of the study. Each subscale is independent of one another and the difference between faculty groups was tested for each subscale separately.

We ran a paired Student’s t-test for course exam metrics comparing FIRST IV faculty to paired comparison faculty. We also ran a simple linear regression of exam metrics against RTOP scores and course size, as the prevalence of scientific practices in assessments may reflect student-centered teaching practices and/or the size of a course.

Sample sizes among the different institution types [86] were insufficient to test for differences in instructional climate (SCII). Despite the uneven distribution of types of institutions, we compared differences in SCII factors (derived from the exploratory factor analysis) across institutions. We performed an ANOVA and Tukey HSD test to compare means of SCII factors for all participating institutions. Additionally, we conducted linear regressions of independent SCII factors with institution size [86] as institution size may affect departmental culture about teaching. To test if departmental climate had differential effects on teaching practices for FIRST IV and comparison faculty, we conducted linear regressions between SCII factors and mean RTOP scores per faculty participant. We tested for differences in the relationship of SCII factors and RTOP between faculty groups with an analysis of covariance (ANCOVA).

### Global model parameter selection

In order to assess the relative importance of external and internal factors on student-centered teaching, we ran a global linear model through model selection using the *dredge* function in the *MuMIn* package in R [87]. This required us to omit faculty participants for whom we did not have SCII institutional data. We performed model selection on a global model with all possible combinations of selected parameters, then ranked models using corrected Akaike’s Information Criterion (AICc) for small sample sizes. We identified the best models as those with a AICc Δ less than two [88]. Once we established the best models, we used model averaging to combine them [89] and derived the relative variable importance from the sum of predicted variable weights and estimates using the full average (model.avg() function in MuMIn package) [88]. All analyses were performed using R version 3.6.1 [90].

### Parameter selection for the global model

Environmental factors that may influence learner-centered teaching practices (RTOP) include engagement in professional development, departmental climate, expectations for faculty, and course constraints (Fig 1). We incorporated professional development into the model as a binary variable indicating whether or not a faculty member had participated in the FIRST IV program. There were significant differences between faculty groups with regards to teaching approach and practice [20], and participation in the FIRST IV program is likely an important variable that affects RTOP scores. Departmental climate, as assessed by the SCII instrument, could influence teaching practices. We opted to use the factor “Mentoring and Material Support” to embody the climate surrounding learner-centered teaching in a department as it had the greatest variation of responses compared to other factors. While there are many requirements of faculty members that may affect teaching practice, we decided to incorporate “Percent Teaching Time Allocation” into the global model. This self-reported perception of the faculty’s job duties reflects their teaching assignment and the time they dedicate to teaching courses as compared to their research and service requirements. Lastly, course constraints are imposed on faculty and may influence their teaching practices. Given the limited course information available, we opted to incorporate average course size per participant into the global model. Course size could influence how an instructor approaches and teaches a course [48, 91].

Internal factors that may influence student-centered teaching practices include instructor self-efficacy and beliefs and intentions surrounding teaching. As each self-efficacy subscale is independent of one another [33], we chose to focus on “Teaching methods” as this subscale is most aligned with student-centered teaching practices in the classroom as assessed by RTOP. There are four independent subscales for the TBI instrument that were generally correlated with one another. We chose to include “Intentions: Knowledge Transmission” in the global model because beliefs don’t always align with practices [92, 93] and there was much higher variation in faculty subscale scores compared to faculty scores for the “Intentions: Learning Facilitation” subscale. This variation may be reflected in teaching practices in the classroom.

### Model selection

The first model selection consisted of determining the relative importance of factors on changes in teaching practice for FIRST IV alumni. The response variable was the change in mean RTOP scores from the end of the FIRST IV program to the present-day study. All variables were scaled to z-scores to account for different ranges across variables. There were 31 faculty that both had four or more RTOP videos [4] and complete SCII scores. The global model included “SCII: Mentoring and Material Support”, “Percent Teaching Appointment”, “Course Size”, “Self-Efficacy: Teaching Methods” and “TBI: Intentions: Knowledge Transmission.”

The second model selection consisted of determining the relative importance of factors on teaching practice for all faculty. The response variable was the mean RTOP score per faculty in the present-day study. All variables were scaled to z-scores to account for different ranges across variables. There were 64 faculty that both had four or more RTOP videos and complete SCII scores. The global model included “FIRST IV faculty/Comparison faculty”, “SCII: Mentoring and Material Support”, “Percent Teaching Appointment”, “Course Size”, “Self-Efficacy: Teaching Methods” and “TBI: Intentions: Knowledge Transmission.”

## Results

### Background Survey

Faculty from both groups considered time and student attitudes/feedback to be the main challenges to implementing active learning in a course (Table 1). The only significant difference between the two groups was with regards to “Classroom infrastructure.” FIRST IV faculty found this obstacle significantly more challenging than comparison faculty (paired Wilcoxon signed-rank test, *P* = 0.026, Cohen’s *d* = 0.485).

**Table 1:**
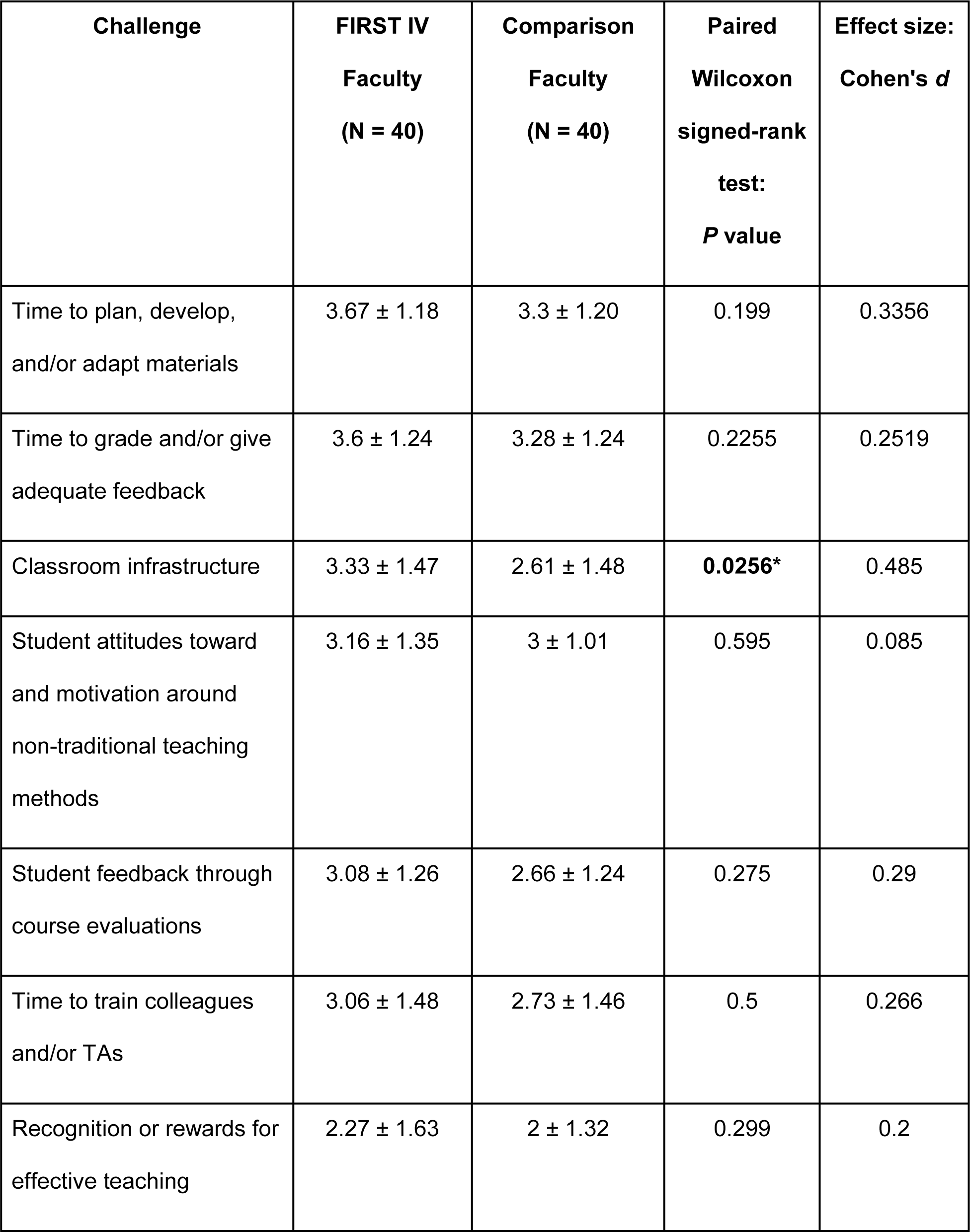
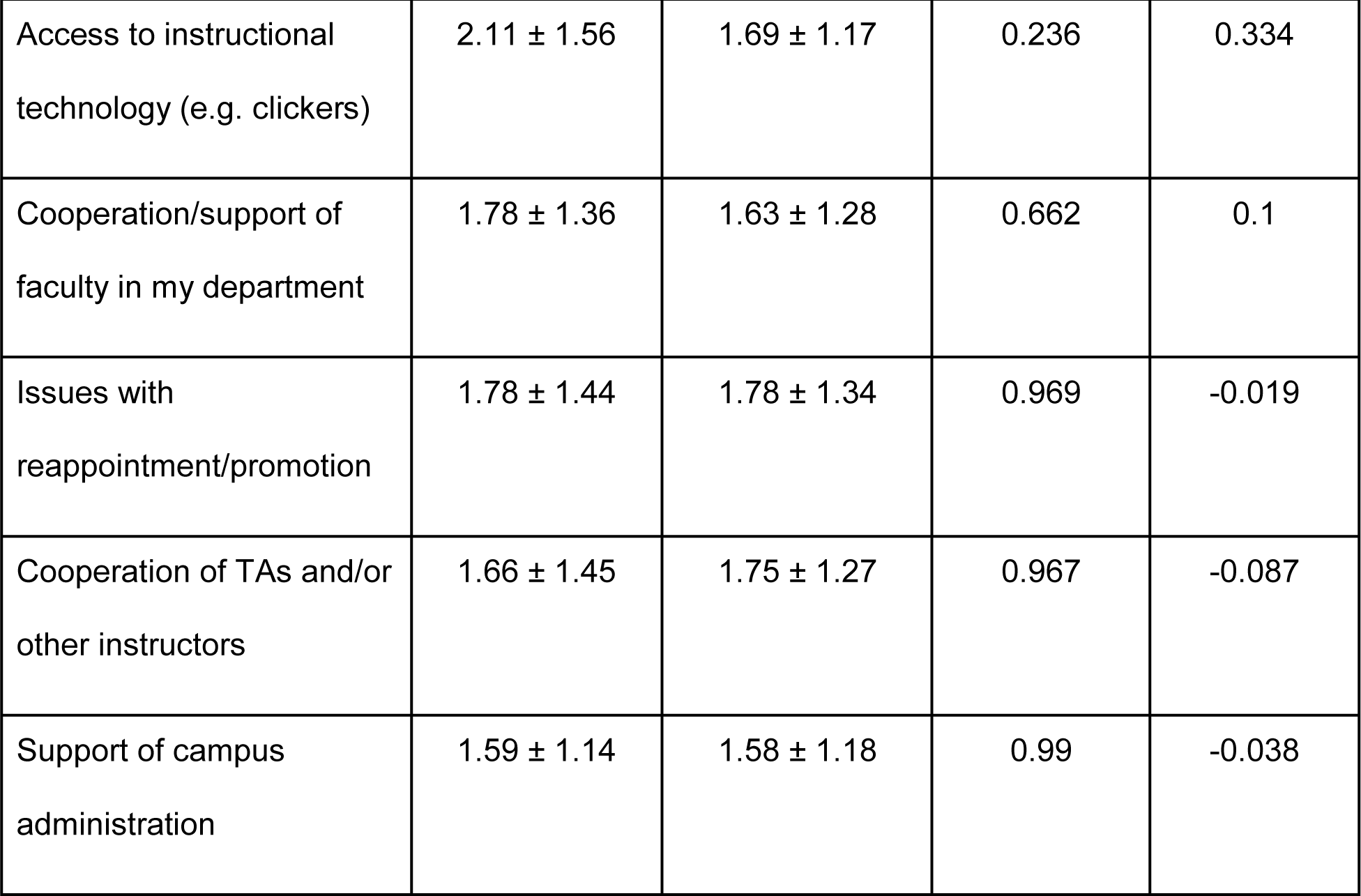
The degree to which the following poses a challenge to implementing active learning in a course (1 = not a challenge, 5 = highly challenging). Reported values for faculty are the mean and standard deviation within a group. Obstacles are sorted by most challenging for FIRST IV faculty, from high to low. Data originate from the background survey.

Faculty groups differed in some regards with respect to their perceptions of departmental culture surrounding teaching (Table 2). Both faculty groups most strongly agreed with the statement: “I frequently discuss issues pertaining to the improvement of teaching and learning with colleagues in my department.” Faculty groups differed significantly for three statements, with comparison faculty significantly agreeing more with each statement than FIRST IV faculty. The statements were: “My department is committed to transforming curricula and courses to enhance active learning and inquiry-based teaching” (paired Wilcoxon signed-rank test, *P* value = 0.0087, Cohen’s *d* = -0.387), “Faculty in my department collaborate to achieve effective teaching (e.g., design, test, discuss curricula, etc.)” (paired Wilcoxon signed-rank test, *P* value = 0.0084, Cohen’s *d* = -0.403), and “Faculty in my department are interested in or are already conducting scholarly work about teaching and learning” (paired Wilcoxon signed-rank test, *P* value < 0.001, Cohen’s *d* = -0.692).

**Table 2:**
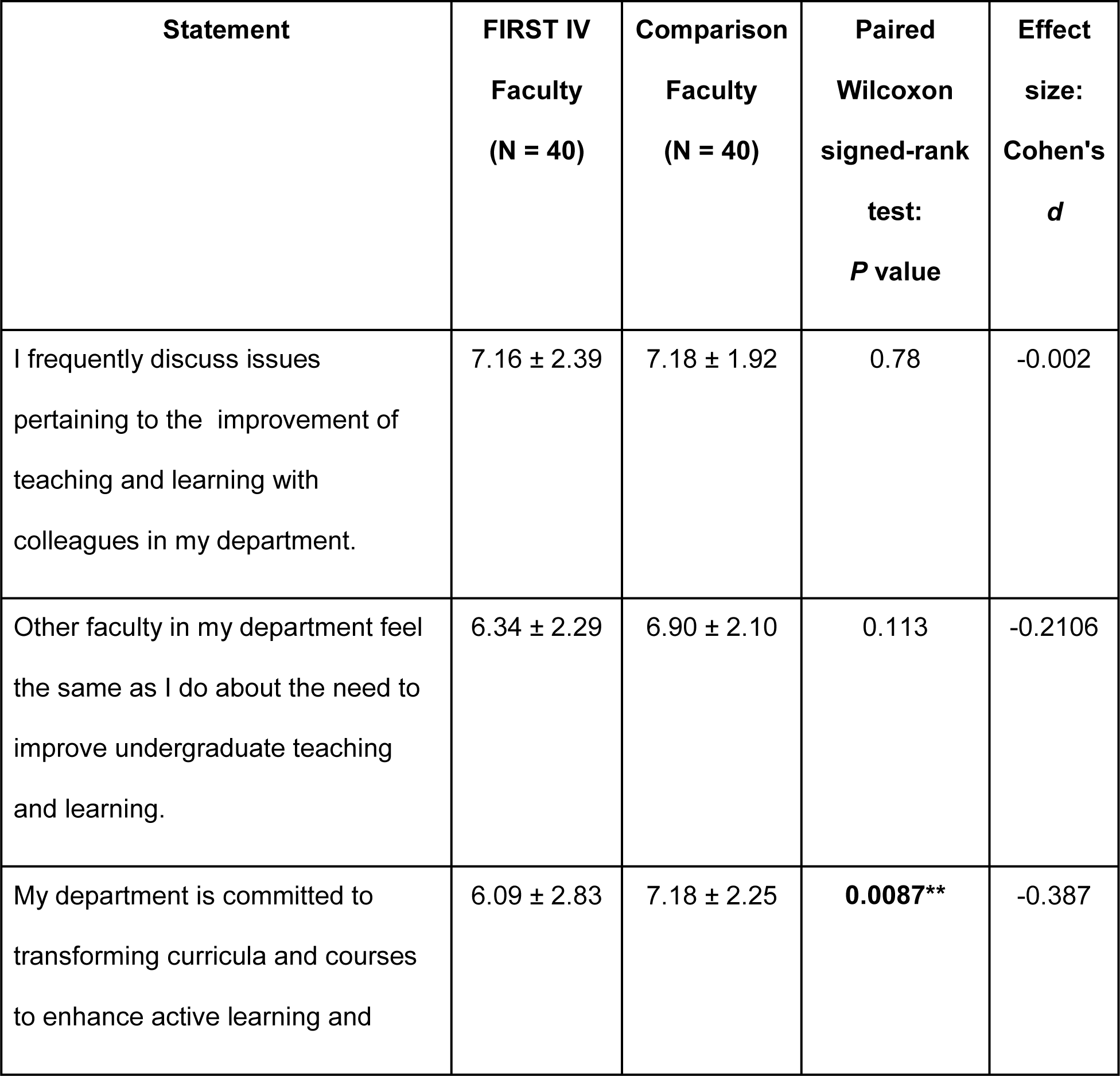
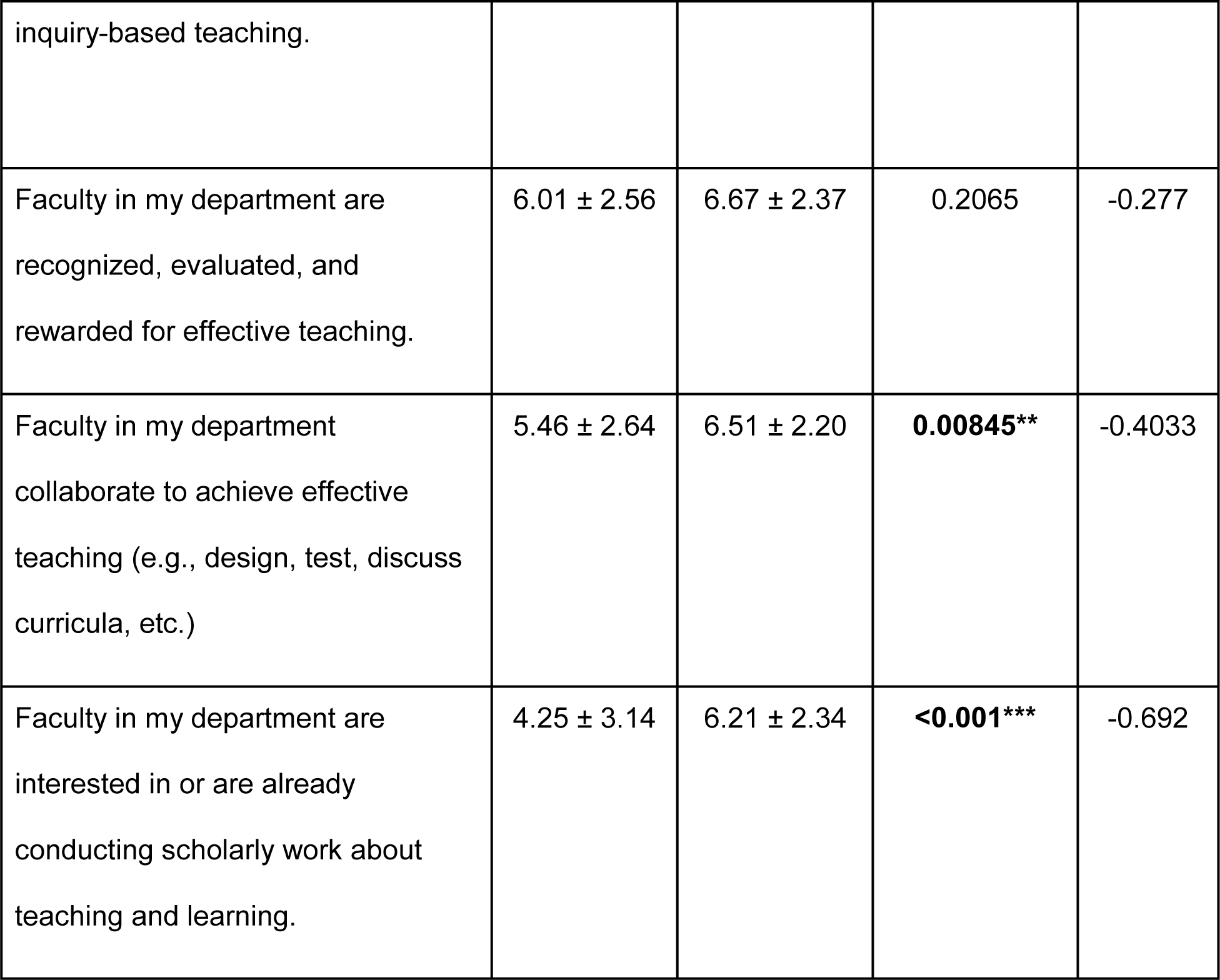
The degree to which faculty agree or disagree with statements about departmental culture around teaching (0 = strongly disagree, 10 = strongly agree). Reported values for faculty are the mean and standard deviation within a group. Statements are ordered by agreement as self-reported by FIRST IV faculty (strongly agree to strongly disagree). Data originate from the background survey.

Differences between faculty groups for self-reported knowledge and experience were consistent across active learning, assessment, and cooperative learning (S3 and S4 Figs). FIRST IV faculty tended to self-report greater knowledge and experience in these three teaching strategies compared to their paired colleagues.

### Teaching Beliefs and Intentions

Faculty had similar teaching beliefs and intentions with regards to “Learning Facilitation” but differed in “Knowledge Transmission” (Fig 2). For “Learning Facilitation” there was no significant difference in Beliefs (paired Wilcoxon signed-rank test, *P* value = 0.878, Cohen’s *d* = 0.034) or Intentions (paired Wilcoxon signed-rank test, *P* value = 0.312, Cohen’s *d* = 0.145). With regards to “Knowledge Transmission,” FIRST IV faculty had significantly lower scores for Beliefs (paired Wilcoxon signed-rank test, *P* value = 0.0082, Cohen’s *d* = -0.599) and Intentions (paired Wilcoxon signed-rank test, *P* value = 0.016, Cohen’s *d* = -0.526).

**Figure 2:**
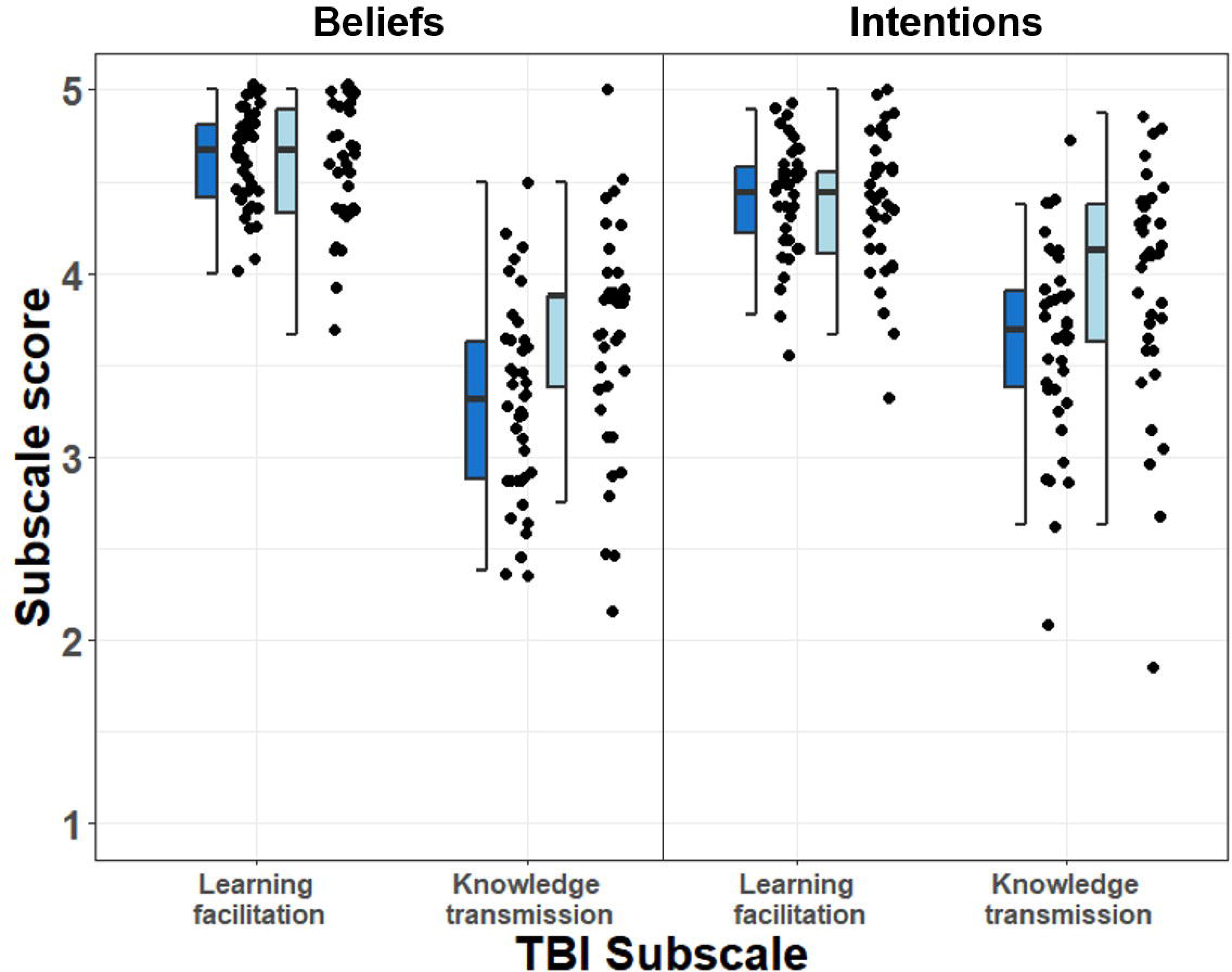
Faculty subscale scores (1-5) for the Teaching Beliefs and Intentions instrument [10]. Dark blue represents FIRST IV faculty (N = 40) and light blue represents comparison faculty (N = 40). The beliefs and intentions of FIRST IV faculty were less likely to align with knowledge transmission teaching practices, as compared to their peers (paired Wilcoxon signed-rank test: Beliefs *P* value = 0.008, Cohen’s *d* = 0.6; Intentions *P* value = 0.016, Cohen’s *d* = 0.53).

### Self-efficacy

Faculty differed in self-efficacy for only one subscale within the self-efficacy instrument (Table 3). FIRST IV faculty had significantly higher self-efficacy than comparison faculty with regards to “Teaching methods” (paired Wilcoxon signed-rank test, *P* value = 0.0088, Cohen’s *d* = 0.561). There were no significant differences between faculty groups for “Course planning,” “Creating learning environment,” “Assessing student learning,” “Interacting with students,” and “Mastering subject knowledge.”

**Table 3:** Faculty subscale scores (1-5) for the Self-Efficacy instrument [33]. Reported values for faculty are the mean and standard deviation within a group.

| **Self-Efficacy**<br>**Subscale** | **FIRST IV**<br>**Faculty**<br>**(N = 40)** | **Comparison**<br>**Faculty**<br>**(N = 40)** | **Paired**<br>**Wilcoxon**<br>**signed-rank**<br>**test:**<br>***P* value** | **Effect**<br>**size:**<br>**Cohen's *d*** |
| --- | --- | --- | --- | --- |
| Course planning | 4.06 ± 0.56 | 3.84 ± 0.6 | 0.089 | 0.389 |
| Teaching methods | 4.16 ± 0.66 | 3.8 ± 0.58 | **0.0088\*** | 0.561 |
| Creating learning<br>environment | 4.09 ± 0.7 | 3.91 ± 0.55 | 0.242 | 0.26 |
| Assessing student learning | 3.96 ± 0.57 | 3.88 ± 0.56 | 0.591 | 0.178 |
| Interact with students | 3.75 ± 0.69 | 3.66 ± 0.45 | 0.716 | 0.14 |
| Mastering subject<br>knowledge | 4.27 ± 0.66 | 4.26 ± 0.56 | 0.745 | -0.041 |

### Departmental Climate

Factor scores for departmental climate varied across institutions and were relatively consistent for “Collegiality”, “Evaluation of Effective Teaching”, “Leadership”, and “Support for Teaching Improvement” (Fig 3). Differences emerged with “Flexibility in Teaching” being significantly higher (*P* <0.001) and “Mentoring and Material Support” being significantly lower than all other factors (*P* <0.001).

**Figure 3:**
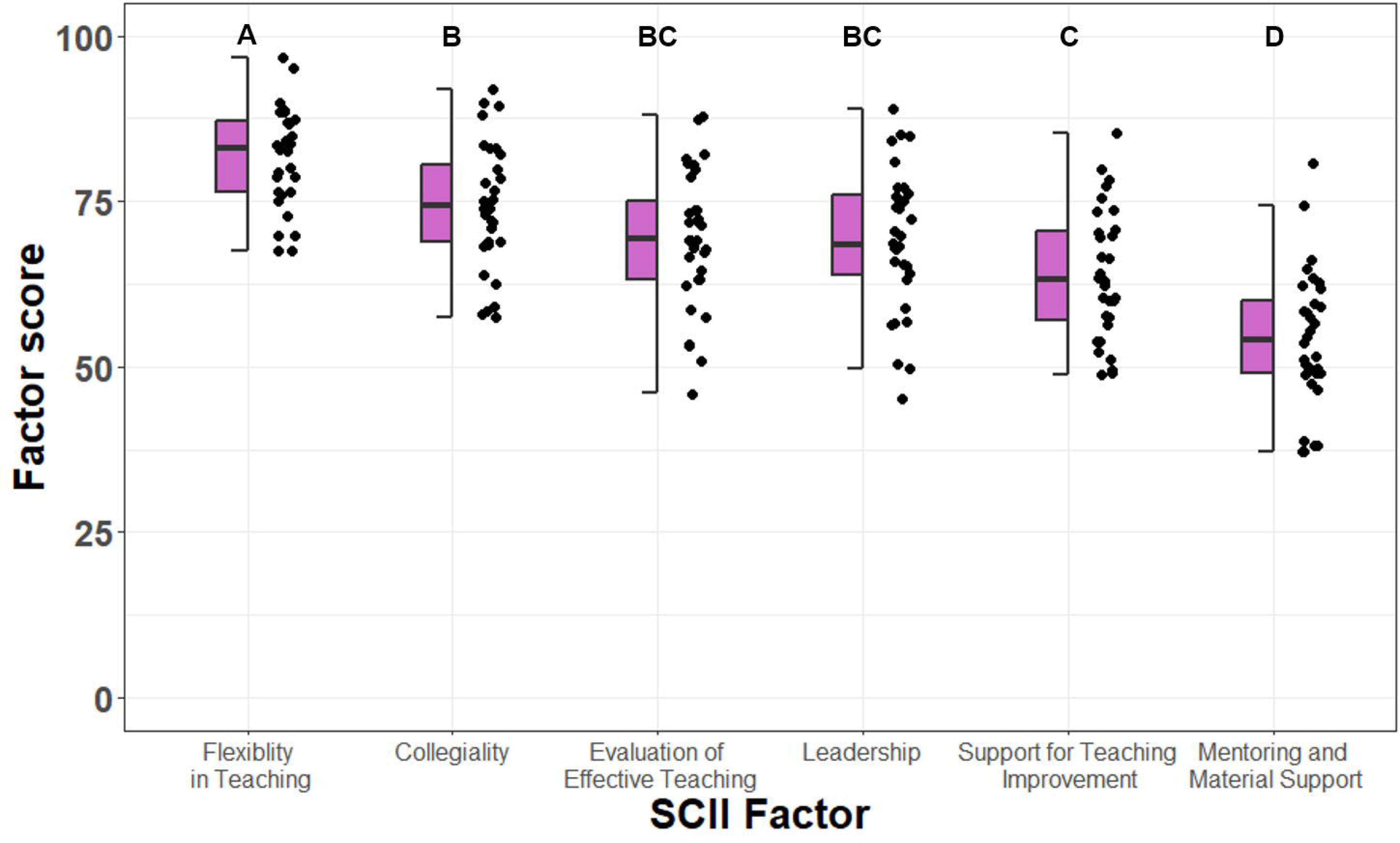
SCII factor scores for all institutions (N = 32). Letters represent significant differences among factors according to an ANOVA and Tukey HSD post hoc test.

When SCII factor scores were regressed by size of institution [86], both “Leadership” and “Evaluation of Effective Teaching” were negatively correlated with institution size (Fig 4; linear regression: A, Leadership, *P* value = 0.044, r^2^ = 0.1; B, Evaluation of Effective Teaching, *P* value = 0.001, r^2^ = 0.27).

**Figure 4:**
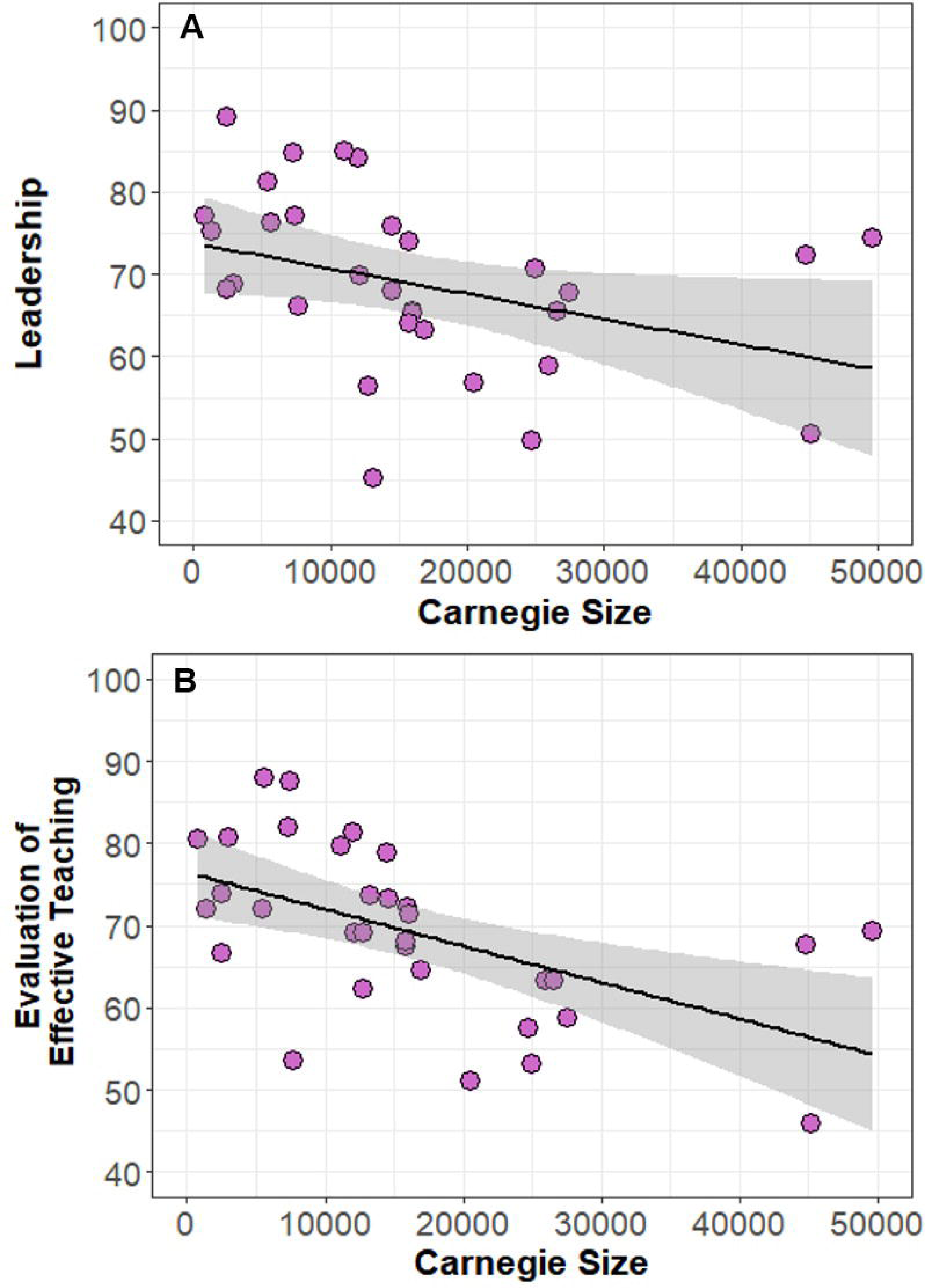
Linear regression for institution size by SCII: Leadership (A; *P* value = 0.044, r^2^ = 0.1) and SCII: Evaluation of Effective Teaching (B; *P* value = 0.001, r^2^ = 0.27). The gray band represents the 95% confidence interval (N = 32 institutions).

All SCII factors had similar relationships with RTOP scores for both faculty groups except for “Collegiality” (Fig 5). While there was no significant relationship for FIRST IV faculty (*P* value = 0.95, r^2^ = 0.0), there was a significant positive correlation for comparison faculty (P value = 0.037, r^2^ = 0.10). The two regression lines were also different from one another (ANCOVA, F =16.31, *P* value = 0.0001).

**Figure 5.**
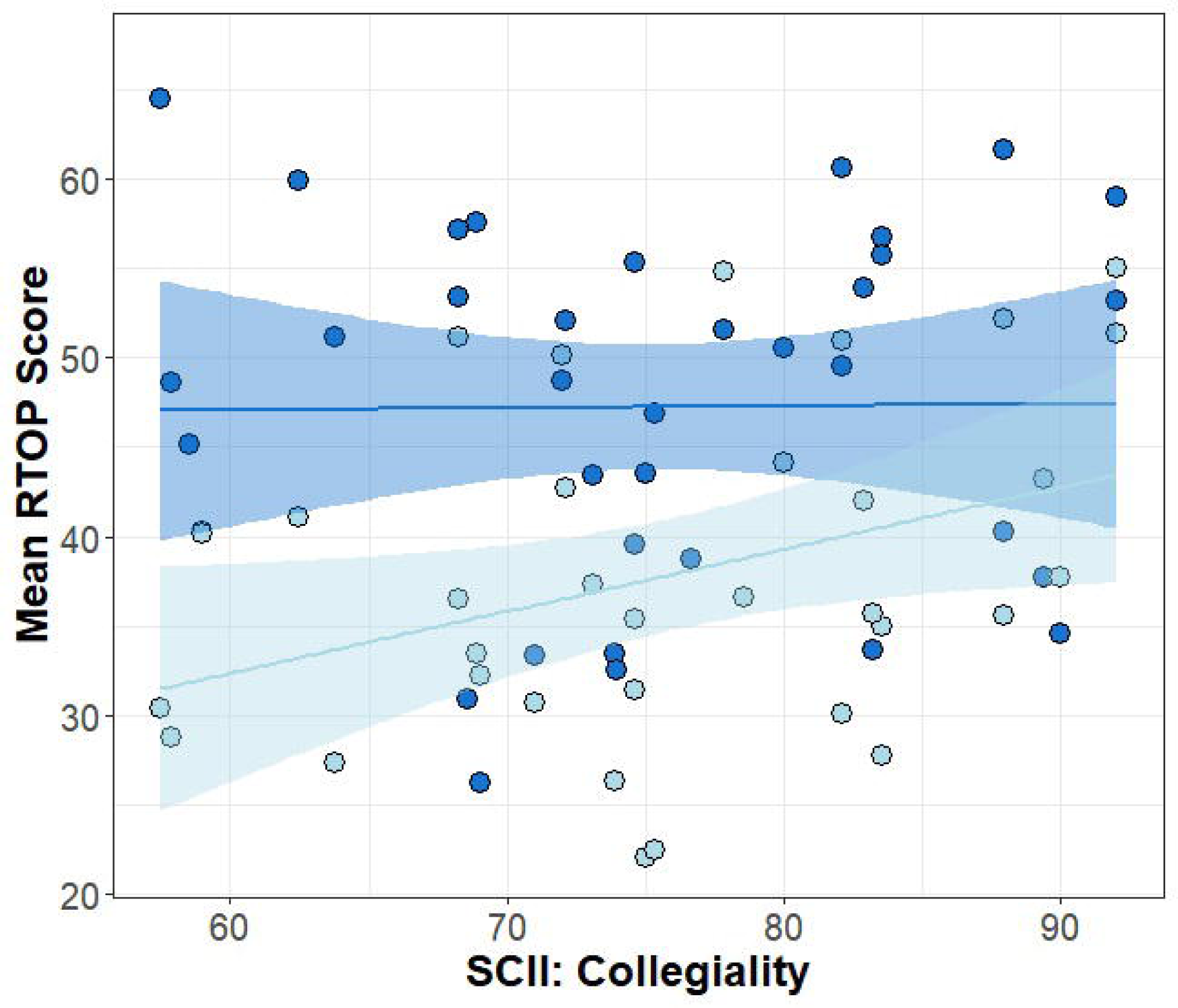
Linear regression of SCII: Collegiality and mean RTOP score for FIRST IV faculty (Dark blue; *P* value = 0.95, r^2^ = 0.0) and comparison faculty (Light blue; *P* value = 0.0001, r^2^ =0.1). The blue bands represent the 95% confidence intervals (N = 70 faculty).

### Model selection

For estimating changes in teaching practice over time, there were six best models with a AICc Δ less than two (S4 Tables). The average of the best models for the change in RTOP scores for FIRST IV alumni (N = 31) consisted of four factors (Table 4). While none of the factors were significant at an alpha = 0.05, the most important factor was “Self-Efficacy: Teaching Methods.” This means that higher confidence in teaching methods leads to a more positive change in RTOP scores over time, and more student-centered teaching practices. The following two factors (in descending order) that comprised the model average were “Intentions: Knowledge Transmission” and “Course Size”, both with negative coefficients. The last factor, with lowest importance, was “Percent Teaching Appointment” with a slight positive coefficient.

**Table 4:**
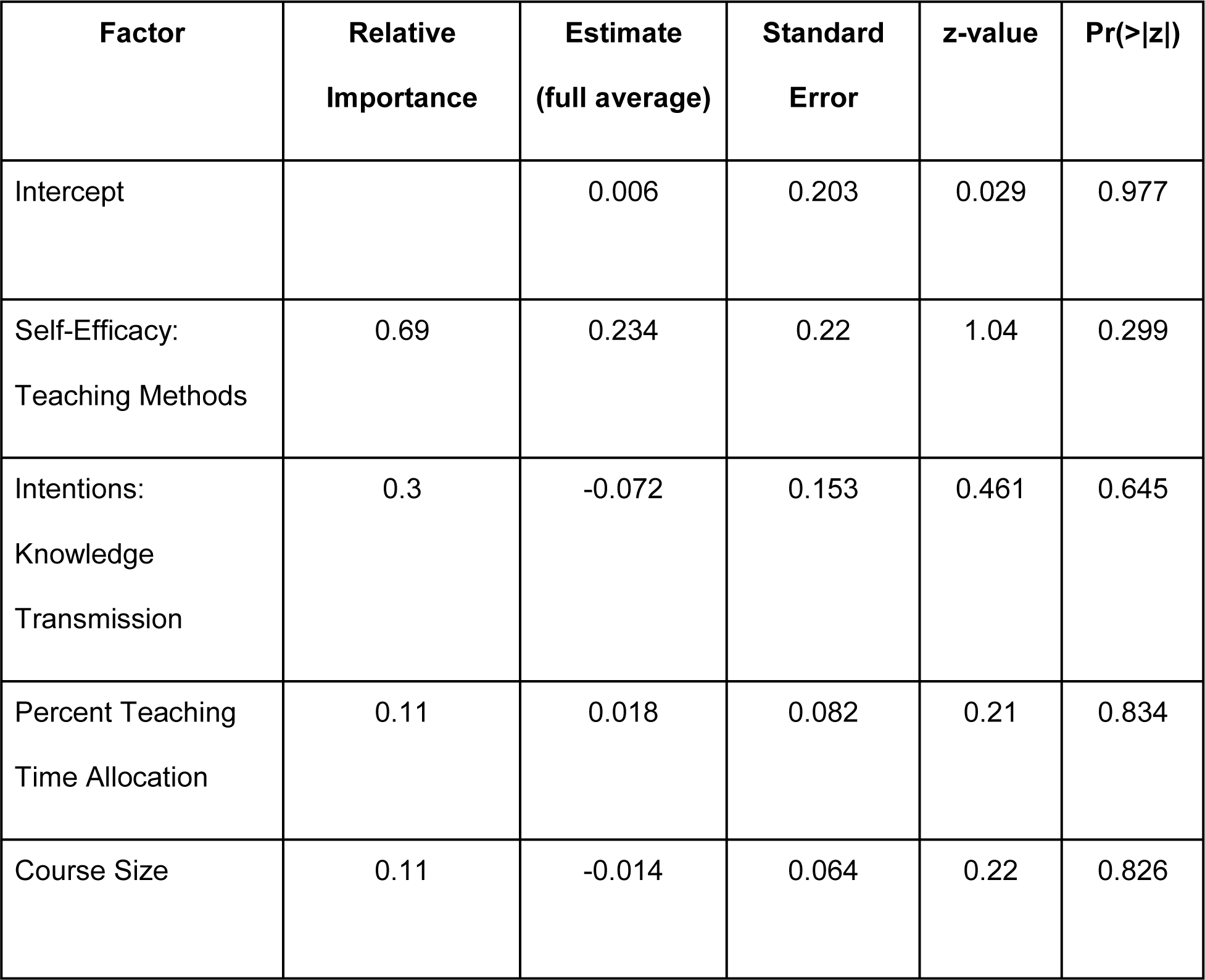
Average model results for the change in RTOP scores for FIRST IV faculty, from the end of FIRST IV to 6-9 years later (N = 31).

| **Factor** | **Relative Importance** | **Estimate (full average)** | **Standard Error** | **z-value** | **Pr(>\|z\|)** |
| --- | --- | --- | --- | --- | --- |
| Intercept |  | 0.006 | 0.203 | 0.029 | 0.977 |
| Self-Efficacy:<br>Teaching Methods | 0.69 | 0.234 | 0.22 | 1.04 | 0.299 |
| Intentions:<br>Knowledge<br>Transmission | 0.3 | -0.072 | 0.153 | 0.461 | 0.645 |
| Percent Teaching<br>Time Allocation | 0.11 | 0.018 | 0.082 | 0.21 | 0.834 |
| Course Size | 0.11 | -0.014 | 0.064 | 0.22 | 0.826 |

For estimating teaching practice across all faculty, there were two best models with a

AICcΔ less than two (S4 Tables). The average of the best models for RTOP scores for all faculty (N =64) consisted of five factors (Table 5). Several of the factors were significant at an alpha = 0.05, including “Self-Efficacy: Teaching Methods”, “Intentions: Knowledge Transmission”, “Percent Teaching Appointment”, and “FIRST IV/Comparison faculty.” With a positive coefficient, higher “Self-Efficacy: Teaching Methods” is related to higher RTOP scores, while greater “Intentions: Knowledge Transmission” and “Percent Teaching Appointment” are related to lower RTOP scores. Not having participated in the FIRST IV program also lowers RTOP scores. The last factor in the average model was “SCII: Mentoring and Material Support”, with a small non-significant positive effect on RTOP scores.

**Table 5:** Average model results for RTOP scores for both FIRST IV and comparison faculty (N =64).

| Factor | Relative Importance | Estimate (full average) | Standard Error | z-value | Pr(>\|z\|) |
| --- | --- | --- | --- | --- | --- |
| Intercept |  | 0.261 | 0.124 | 2.068 | 0.039 |
| Self-Efficacy: Teaching Methods | 1 | 0.442 | 0.092 | 4.72 | **<0.001\*\*\*** |
| FIRST IV Professional Development | 1 | -0.425 | 0.193 | 2.16 | **0.031\*** |
| Intentions:<br>Knowledge<br>Transmission | 1 | -0.350 | 0.098 | 3.50 | **<0.001\*\*\*** |
| Percent Teaching<br>Time Allocation | 1 | -0.203 | 0.092 | 2.17 | **0.03\*** |
| SCII: Mentoring and<br>Material Support | 0.62 | 0.098 | 0.103 | 0.941 | 0.346 |

## Discussion

The way that faculty teach is shaped by both their individual characteristics and environmental influences. Professional development experiences, departmental climate, time allocation, and course type can all impact how faculty instruct. Additionally, faculty themselves hold beliefs and intentions about teaching and may have different levels of confidence in their ability to teach a learner-centered course. Our study found that self-efficacy in teaching exerts a strong influence on faculty teaching practices (Table 5) and potentially influences how those practices may change over time (Table 4). Additionally, faculty intentions about teaching in the classroom also appear to significantly influence their practice. Environmental influences play a role in supporting or constraining faculty teaching but ultimately, individual characteristics appear to have a greater role in shaping teaching practices. Among those environmental influences, participation in professional development, in this case the FIRST IV program, appears to have a significant effect on teaching practices.

### Self-efficacy in teaching is important for all faculty

Perceived self-efficacy [28 pg. 3] is the “belief in one’s capabilities to organize and execute the courses of action required to produce given attainments.” Faculty are responsible for numerous tasks and skills in academia, and thus may have various degrees of self-efficacy among their work obligations [30, 32]. Our results suggest that self-efficacy in teaching is a significant influence on learner-centered teaching practices (Table 5). Additionally, changes in FIRST IV faculty teaching practices over time seem to be positively influenced by self-efficacy in teaching (Table 4). As faculty confidence in learner-centered teaching methods increased, so did the degree of learner-centered teaching practices used in the classroom. This is consistent with previous findings relating the importance of instructor self-efficacy to teaching practices [31,34,94,95]. Self-efficacy is attributed to many sources, including mastery experiences, social persuasions, and professional development [9,35,33,96]. Confidence in teaching methods possibly stems from knowledge and experience gained during the FIRST IV program. FIRST IV faculty tended to report greater knowledge and experience in learner-centered teaching than comparison faculty (S3 and S4 Figs). The difference in self-efficacy between faculty groups may be a reflection of the training that faculty experienced during the FIRST IV professional development program. Graduate programs and training for early-career academics could have tremendous benefits for faculty teaching self-efficacy that result in shifting teaching practices in the classroom [33,34,97,98]. Our results suggest that developing confidence and self-efficacy in teaching for early-career academics through professional development, such as FIRST IV, could increase the prevalence of learner-centered teaching practices in higher education.

### FIRST IV program makes a difference

The FIRST IV professional development program resulted in long-term impacts for early-career faculty [20]. Our results suggest that not only do FIRST IV faculty approach and teach courses from a more learner-centered perspective than their peers [20], but also they are also less supportive of knowledge transmission in teaching settings (Fig 2). Previous research has also found that professional development can lead to lower intentions towards knowledge transmission [99]. In addition to differences in teaching beliefs and intentions, FIRST IV faculty self-reported greater self-efficacy in teaching methods (Table 3). These results reinforce the conclusion that this professional development program, which worked with participants when they were postdocs, has had long-term outcomes as they transitioned to faculty positions at institutions across the country. The FIRST IV program was likely effective at shaping teaching approach, practice, and confidence with regards to student-centered learning. These results are supported by the importance of the FIRST IV program in the model of learner-centered teaching practice (Table 5). Participation in FIRST IV was a significant predictor of RTOP scores, reaffirming the value of teaching professional development in improving scientific teaching practices in the classroom [18,100,101].

Teaching professional development not only seeks to change participant attitudes and behavior, but also can affect how participants view the work environment [102, 103]. In our study, FIRST IV faculty perceived departmental climate for teaching differently than their peers. While both faculty groups felt similarly about communication among instructors, differences emerged with FIRST IV faculty disagreeing more with statements about a department’s commitment to change, collaboration among faculty, and the interest in or extent of scholarship in teaching and learning conducted in their department (Table 2). Compared to their peers, FIRST IV faculty do not consider their department and colleagues to be as committed to transforming courses or as open to collaboration. The significant differences in the perception of departmental culture may faculty (S3 and S4 Figs), since the FIRST IV program was highly collaborative and explored scholarship in teaching and learning in depth. Discussing pedagogy and working together with other instructors can have tremendous benefits for faculty looking to change their teaching approaches and practices [102,104–106]. With that exposure, FIRST IV faculty possibly became more aware of what it means to effectively transform curricula, collaborate, and conduct scholarship in teaching and learning. It is also possible that FIRST IV faculty built a teaching-focused social network that continues to provide insight and perspective on academic culture surrounding teaching [107]. By changing individual faculty perceptions and attitudes in a collaborative environment, it may be possible to affect long-term cultural change surrounding teaching practices. This increased awareness is yet another potential benefit of teaching professional development programs for early-career academics.

### Environmental constraints have some impact

Faculty teaching is often constrained by time and course characteristics [27, 108]. Participants in our study agree that time and student feedback were challenges to implementing active learning in the classroom (Table 1). Having sufficient time to develop and implement active learning exercises may depend on one’s time allocation to teaching. Time allocation can hinder faculty self-reflection with respect to teaching practices and curb their growth as an instructor [109]. Teaching-focused faculty might have more time to develop and implement learner-centered teaching practices than research-focused faculty. However, in our model of teaching practice, faculty time allocation to teaching was a significant predictor of learner-centered teaching practices with a negative estimate (Table 5). This suggests that the greater a faculty’s percent teaching responsibilities, the lower their RTOP score. Percent time allocated to teaching was self-reported. Conflict between the amount of time dedicated to teaching and the focus on research scholarship for promotion and tenure is well established [110, 111]. Faculty who are evaluated on research productivity and have a high number of teaching responsibilities may not feel like they have time to teach learner-centered courses. Alternatively, teaching-focused faculty may be overwhelmed with teaching responsibilities [46] and not have sufficient time to dedicate to learner-centered practices, as our study participants cited time as the greatest challenge to implementing active learning in the classroom (Table 1). How and why faculty allocate their time and energy is a complex story and depends on many factors. However, developing and implementing scientific teaching practices in the classroom does require time in a faculty’s schedule. If departments and faculty leaders are committed to transforming courses and curricula to be more learner-centered, they need to recognize the value of providing time for incorporating feedback [112], and course development [113].

A notable course constraint observed in our study was classroom infrastructure. FIRST IV faculty felt strongly that classroom infrastructure was a challenge to implementing active learning in the classroom to a greater extent than their peers (Table 1). This significant difference in faculty perceptions of how classroom structure affects active learning may stem from FIRST IV alumni knowledge of and experience with spaces designed for learner-centered teaching (S3 and S4 Figs). Knowledge of active learning and how to implement it in different types of classrooms takes training and practice, an element that differs significantly between FIRST IV faculty and comparison faculty [20]. In addition to classroom infrastructure, course enrollment possibly affects teaching practices [48, 91]. However, in our study, course size was not a significant factor in the overall model of teaching practice, and had a very small contribution to our model of RTOP scores over time for FIRST IV faculty. Past research has also found that small course enrollment does not necessarily result in the implementation of learner-centered teaching [4,69,114]. Overall, course infrastructure was more of a perceived barrier for faculty trained in scientific teaching and course size does not seem to have a significant effect on learner-centered teaching practices.

### Departmental climate has a limited role

Faculty teach courses as part of a curriculum and work with colleagues on course development on many levels, including goals, instructional designs and assessments. Thus, it is not unreasonable to hypothesize that departmental climate has an effect on actual instruction [69]. Institutional and departmental climate can be a barrier to implementing research-based teaching techniques [58]. By building a community around teaching and facilitating collegial interactions and collaboration, departments can improve teaching practices [115–118], which is especially important for early-career faculty [71]. Additionally, supportive leadership and a positive teaching climate can influence student perceptions and experiences in the classroom [72, 73]. In our model of learner-centered teaching practices, “Mentoring and Material Support” had a non-significant positive effect on teaching practices in the classroom (Table 5), but for FIRST IV faculty, there was no effect of departmental climate on changing faculty teaching practices over time (RTOP). While past literature has suggested and found some evidence of departmental climate on teaching practices, in our model of faculty teaching practice, climate does not appear to have as important a role as individual faculty characteristics.

Departmental climate can vary across institutions and we found that larger institutions tended to have lower factor scores for “Leadership” and “Evaluation of Effective Teaching” (Fig 4). At larger institutions, departments may be more research-focused and less focused on shifting to student-centered learning environments. The leadership in these departments may also promote a culture of focusing on research more than teaching. Additionally, while “Flexibility in Teaching” and “Collegiality” were high across all institutions, “Mentoring and Material Support” was significantly lower than other climate characteristics (Fig 3). This result provides evidence for the need for greater mentorship and sustained support for instructors to teach learner-centered courses. With respect to flexibility and collegiality, it is encouraging that faculty encountered positive interactions with their peers about teaching and flexibility in their teaching approaches across a variety of institutions. Peer interactions can be important for changing teaching practices in higher education [12]. This is also evident from the relationship between collegiality and teaching practices between the two groups of faculty (Fig 5). While FIRST IV faculty teaching is unaffected by differences in collegiality, a more collegial departmental climate had a positive effect on teaching practices for faculty who had not participated in significant professional development. Thus, trained faculty appear to be more resilient to unfavorable teaching climates than untrained faculty. When faculty arrive at a new institution and a new community, they may struggle to incorporate research-based teaching practices. Peer-peer interactions and conversations around teaching can help early-career faculty become better instructors [119, 120]. As postdocs, the FIRST IV faculty established a community of skilled, motivated instructors and mentors that continues to this day through an active private email listserv. It is possible that professional development from the FIRST IV program and the supportive community that was established has enabled these instructors to teach learner-centered courses in a variety of collegial teaching climates. In addition to improving instructor approaches and practices, teaching professional development may result in confident, resilient teachers that can engage students in various departmental climates. Ultimately, transforming departmental climate surrounding teaching can bring about change in higher education practices [121]. Meaningful change in higher education teaching practices should come from administration as well as instructors. Although our results suggest that the individual is more important for teaching practices than the environment, departmental climate and institutional support still play a critical role in the nurturing and encouraging of learner-centered teaching approaches and practices.

### Limitations

While this study sought to investigate questions that apply to early-career biology faculty, the distribution of participating institutions was uneven. The majority of participants came from doctoral and master’s granting institutions (31 out of 35) with only two baccalaureate and two community colleges represented. The results are thus more likely to reflect the importance of different factors on learner-centered teaching for faculty at larger institutions that have a greater research focus. Additionally, FIRST IV alumni applied to and accepted offers of employment from the departments in our study. It is possible that these departments are generally more supportive of learner-centered teaching than departments not involved in this study.

While our results have broad implications for facilitating learner-centered teaching practices across STEM disciplines, this study was conducted with primarily biology faculty and there are known disciplinary differences in teaching approaches and practices in higher education [69]. However, while our study participants were mainly biologists, the disciplinary scope of each department varied from specific to broad. We believe that our study incorporates much of the variation observed in department support for learner-centered teaching practices.

Finally, each of the factors influencing teaching practice do not act independently. They are inherently interacting with one another, for example, professional development can affect self- efficacy in teaching [38]. Due to the limited number of faculty participants in our study we were unable to incorporate these potential interactions. Although much of the data are derived from published self-reported instruments, our response variable is derived from direct observations of teaching as scored by RTOP. Despite limitations, we are confident that our models contain critical variables or characteristics that affect teaching practices and reveal their relative importance.

## Conclusions

Faculty teaching practices in the classroom are the result of both the instructor characteristics and their surrounding environment. Our study found that the individual characteristics, in particular self-efficacy, were most important in determining the degree to which a classroom was learner-centered. In general, environmental factors had a limited effect on teaching practices, although professional development had a significant positive effect on teaching. Departmental climate did not emerge as a significant factor in our model, but there was a consistent pattern across all the departments in our study, that mentoring and material support for teaching was low and that aspects of the instructional climate for teaching were negatively related to institution size. While faculty perceived departments’ lack of mentoring and material support for teaching, many trained instructors persevere with confidence in their abilities and an active teaching support network. Our results suggest that through professional development and cultivating instructor self-efficacy, STEM teaching practices can effectively shift towards a learner-centered approach.

## Author contributions

NE collected data, analyzed the results, and wrote the manuscript.

JMM designed the study, collected data, wrote and edited the manuscript. DE-M designed the study, collected data, wrote and edited the manuscript.

## Acknowledgements

The NSF funded this study (DUE-1623834) for which we are most grateful. We thank our advisory committee (Sara Brownell, Matt Hora, Charles Henderson, Mark Connolly, and Kevin Ford) for their insight on data collection and analysis of the results. We appreciate the work of Sarah Eddy for statistical consultation on the model. An especially big thanks to all of the participating faculty and students who comprised the data collected in this study.

## Supporting information

**S1. Relevant Background Survey questions.**

**S2. SCII items and exploratory factor analysis loadings for a six factor structure.**

**S3 Fig. Knowledge about teaching strategies.** These are faculty responding to their knowledge level with respect to active learning, assessment, and cooperative learning. The percentage is the percent of faculty reporting a certain level of knowledge within a group (FIRST IV or comparison).

**S4 Fig. Experience about teaching strategies.** These are faculty responding to their experience level with respect to active learning, assessment, and cooperative learning. The percentage is the percent of faculty reporting a certain level of experience within a group (FIRST IV or comparison).

**S5 Tables**. AICc tables for DeltaRTOP & RTOP overall models

**S6**. Data column headers

## References

1. National Research Council. Discipline-based education research: understanding and improving learning in undergraduate science and engineering. National Academies Press, Washington, DC. 2012.

2. National Academies of Sciences, Engineering, and Medicine. Indicators for monitoring undergraduate STEM education. Washington, DC: The National Academies Press. 2018.

3. Freeman S, Eddy SL, McDonough M, Smith MK, Okoroafor N, Jordt H, Wenderoth MP. Active learning increases student performance in science, engineering, and mathematics. Proceedings of the National Academy of Sciences, 2014;111: 8410–8415.

4. Stains M, Harshman J, Barker MK, Chasteen SV, Cole R, Dechenne-Peters SE, Eagan MK, Esson JM, Knight JK, Laski FA, Levis-Fitzgerald M. Anatomy of STEM Teaching in American Universities: A Snapshot from a Large-Scale Observation Study. Science, 2018;359: 1468–1470.

5. Theobald EJ, Hill MJ, Tran E, Agrawal S, Arroyo EN, Behling S, Chambwe N, Cintrón DL, Cooper JD, Dunster G, Grummer JA. Active learning narrows achievement gaps for underrepresented students in undergraduate science, technology, engineering, and math. Proceedings of the National Academy of Sciences. 2020;117: 6476–83.

6. Cromley JG, Perez T, Kaplan A. Undergraduate STEM achievement and retention: Cognitive, motivational, and institutional factors and solutions. Policy Insights from the Behavioral and Brain Sciences. 2016;3:4–11.

7. Jordt H, Eddy SL, Brazil R, Lau I, Mann C, Brownell SE, King K, Freeman S. Values affirmation intervention reduces achievement gap between underrepresented minority and white students in introductory biology classes. CBE—Life Sciences Education. 2017;16.

8. Madson L, Trafimow D, Gray T. Faculty Members’ Attitudes Predict Adoption of Interactive Engagement Methods. The Journal of Faculty Development, 2017;31: 39–50.

9. Bray-Clark N, Bates R. Self-efficacy beliefs and teacher effectiveness: Implications for professional development. Professional Educator, 2003;26: 13–22.

10. Norton L, Richardson TE, Hartley J, Newstead S, Mayes J. Teachers’ beliefs and intentions concerning teaching in higher education. Higher Education, 2005;50: 537–571.

11. Andrews TC, Lemons PP. It’s personal: Biology instructors prioritize personal evidence over empirical evidence in teaching decisions. CBE Life Sciences Education, 2015;14: 1–18.

12. Auerbach AJ, Schussler E. A vision and change reform of introductory biology shifts faculty perceptions and use of active learning. CBE Life Sciences Education, 2017;16: 1–12.

13. Derting TL, Ebert-May D, Henkel TP, Maher JM, Arnold B, Passmore HA. Assessing faculty professional development in STEM higher education: Sustainability of outcomes. Science Advances, 2016;2.

14. Austin AE. Institutional and departmental cultures: The relationship between teaching and research. New Directions for Institutional Research, 1996;90: 57–66.

15. Miller ER, Fairweather JS. The Role of Cultural Change in Large-Scale STEM Reform: The Experience of the AAU Undergraduate STEM Education Initiative. Transforming Institutions: Undergraduate STEM Education for the 21st Century, 2015;48.

16. Henderson C, Beach A, Finkelstein N. Facilitating change in undergraduate STEM instructional practices: An analytic review of the literature. Journal of Research in Science Teaching, 2011;48: 952–984.

17. Teasdale R, Viskupic K, Bartley JK, McConnell D, Manduca C, Bruckner M, … Iverson E. A multidimensional assessment of reformed teaching practice in geoscience classrooms. Geosphere, 2017;13: 608–627.

18. Viskupic K, Ryker K, Teasdale R, Manduca C, Iverson E, Farthing D, … McFadden R. Classroom Observations Indicate the Positive Impacts of Discipline-Based Professional Development. Journal for STEM Education Research, 2019;2: 201–228.

19. Emery N, Maher JM, Ebert-May D. Studying Professional Development as Part of the Complex Ecosystem of STEM Higher Education. Innovative Higher Education, 2019;44: 469–479.

20. Emery N, Maher JM, Ebert-May D. Early-career faculty practice learner-centered teaching up to 9 years after postdoctoral professional development. Science Advances, 2020;6

21. Stes A, Coertjens L, Van Petegem P. Instructional development for teachers in higher education: impact on teaching approach. Higher Education, 2010;60: 187–204.

22. Stains M, Pilarz M, Chakraverty D. Short and Long-Term Impacts of the Cottrell Scholars Collaborative New Faculty Workshop. Journal of Chemical Education, 2015;92: 1466–1476.

23. Manduca CA, Iverson ER, Luxenberg M, Macdonald RH, McConnell DA, Mogk DW, Tewksbury BJ. Improving undergraduate STEM education: The efficacy of discipline-based professional development. Science Advances, 2017;3: 1–16.

24. Bandura A. Social foundations of thought and action: A social cognitive theory. Englewood Cliffs, NJ: Prentice Hall. 1986.

25. Oleson A, Hora MT. Teaching the way they were taught? Revisiting the sources of teaching knowledge and the role of prior experience in shaping faculty teaching practices. Higher education, 2014;68: 29–45.

26. Azjen I. The theory of planned behavior. Organizational Behavior and Human Decision Processes, 1991;50: 179–211.

27. Brownell SE, Tanner KD. Barriers to faculty pedagogical change: Lack of training, time, incentives, and tensions with professional identity? CBE Life Sciences Education, 2012;11: 339–346.

28. Bandura, A. Self-efficacy: The exercise of control. Macmillan. 1997.

29. Scholz U, Doña BG, Sud S, Schwarzer R. Is general self-efficacy a universal construct? Psychometric findings from 25 countries. European Journal of Psychological Assessment, 2002;18: 242–251.

30. Landino RA, Owen SV. Self-efficacy in university faculty. Journal of Vocational Behavior, 1988;33: 1–14.

31. Pajares F. Self-efficacy beliefs in academic settings. Review of Educational Research, 1996;66: 543–578.

32. Bailey JG. Academics’ motivation and self efficacy for teaching and research. Higher D Education Research & Development, 1999;18: 343–359.

33. Connolly MR, Lee YG, Savoy JN. The effects of doctoral teaching development on early-career STEM scholars’ college teaching self-efficacy. CBE—Life Sciences Education, 2018;17: ar14.

34. Fong CJ, Dillard JB, Hatcher M. Teaching self-efficacy of graduate student instructors: Exploring faculty motivation, perceptions of autonomy support, and undergraduate student engagement. International Journal of Educational Research, 2019;98: 91–105.

35. Morris DB, Usher EL. Developing teaching self-efficacy in research institutions: A study of award-winning professors. Contemporary Educational Psychology, 2011;36: 232–245.

36. Morris DB, Usher EL, Chen JA. Reconceptualizing the sources of teaching self-efficacy: A critical review of emerging literature. Educational Psychology Review, 2017;29: 795–833.

37. Singh T, de Grave W, Ganjiwale J, Supe A, Burdick WP, van der Vleuten C. Impact of a fellowship program for faculty development on the self-efficacy beliefs of health professions teachers: a longitudinal study. Medical teacher, 2013;35: 359–364.

38. Rowbotham MA. The Impact of Faculty Development on Teacher Self-Efficacy, Skills and Perspectives. Policy Research: IERC FFR 2015-1. Illinois Education Research Council. 2015.

39. Sitzmann T, Ely K. A meta-analysis of self-regulated learning in work-related training and educational attainment: What we know and where we need to go. Psychological bulletin,2011;137: 421.

40. Milem JF, Berger JB, Dey, EL. Faculty time allocation: A study of change over twenty years. The Journal of Higher Education, 2000;71: 454–475.

41. Umbach PD. The effects of part-time faculty appointments on instructional techniques and commitment to teaching. In 33rd annual conference of the association for the study of higher education, Jacksonville, FL (Vol. 58). 2008.

42. Baldwin RG, Wawrzynski MR. Contingent faculty as teachers: What we know; what we need to know. American Behavioral Scientist, 2011;55: 1485–1509.

43. Figlio DN, Schapiro MO, Soter KB. Are tenure track professors better teachers? Review of Economics and Statistics, 2015;97: 715–724.

44. Rawn CD, Fox JA. Understanding the work and perceptions of teaching focused faculty in a changing academic landscape. Research in Higher Education, 2018;59: 591–622.

45. Bush SD, Rudd JA, Stevens MT, Tanner KD, Williams KS. Fostering change from within: Influencing teaching practices of departmental colleagues by science faculty with education specialties. PLoS ONE, 2016;11: 1–20.

46. Xu D, Solanki S. Tenure-Track Appointment for Teaching-Oriented Faculty? The Impact of Teaching and Research Faculty on Student Outcomes. Educational Evaluation and Policy Analysis, 2020;42: 66–86.

47. Braxton JM, Berger JB. Public trust, research activity, and the ideal of service to students as clients of teaching. New Directions for Institutional Research, 1996;90: 79–91.

48. Cuseo J. The empirical case against large class size: Adverse effects on the teaching, learning, and retention of first-year students. The Journal of Faculty Development, 2007;21: 5–21.

49. Ratcliff JL. What they took and what they learned: Learning from assessment and transcript analysis. In M. Moseley (Ed.), Proceedings from the Asheville Institute on General Education (pp. 64-69). (A program of the Association of American Colleges and The University of North Carolina-Asheville) Washington, D.C.: Association of American Colleges. 1992.

50. Carbone E, Greenberg J. Teaching large classes: Unpacking the problem and responding creatively. In M. Kaplan (Ed.), To improve the academy, vol. 17, Stillwater, OK: New Forums Press and The Professional and Organizational Development Network in Higher Education. 1998.

51. Mulryan-Kyne C. Teaching large classes at college and university level: challenges and opportunities. Teaching in Higher Education, 2010;15: 175–185.

52. Hamilton WR, Padron VA, Henriksen JA. Engaging Students in a Large Classroom and Distance Environment. In Handbook of Research on Teaching and Learning in K-20 Education (pp. 759–777). IGI Global. 2013.

53. Hornsby DJ, Osman R. Massification in higher education: Large classes and student learning. Higher education, 2014;67: 711–719.

54. Dean T, Lee-Post A, Hapke H. Universal design for learning in teaching large lecture classes. Journal of Marketing Education, 2017;39: 5–16.

55. Henshaw RG, Edwards PM, Bagley EJ. Use of Swivel Desks and Aisle Space to Promote Interaction in Mid-Sized College Classrooms. Journal of Learning Spaces, 2001;1: n1.

56. Casanova D, Di Napoli R, Leijon M. Which space? Whose space? An experience in involving students and teachers in space design. Teaching in Higher Education, 2018;23: 488–503.

57. Chen PSD, Lambert AD, Guidry KR. Engaging online learners: The impact of Web-based learning technology on college student engagement. Computers & Education, 2010;54: 1222–1232.

58. Henderson C, Dancy MH. Barriers to the use of research-based instructional strategies: The influence of both individual and situational characteristics. Physical Review Special Topics-Physics Education Research, 2007;3

59. Postareff L, Lindblom-Ylänne S, Nevgi A. The effect of pedagogical training on teaching in higher education. Teaching and teacher education, 2007;23: 557–571.

60. Pfund C, Miller S, Brenner K, Bruns P, Chang A, Ebert-May D, Fagen AP, Gentile J, Gossens S, Khan IM, Labov J. Summer Institute to Improve University Science Teaching. Science, 2009;324: 470–471.

61. Stes A, Clement M, Van Petegem P. The Effectiveness of a Faculty Training Programme: Long-term and institutional impact. International Journal for Academic Development, 2007;12: 99–109.

62. Tennill MM, Cohen MW. 9: Assessing the long-term impact of a professional development program. To Improve the Academy. 2013;32: 145–59.

63. Stewart M. Making sense of a teaching programme for university academics: Exploring the longer-term effects. Teaching and Teacher Education. 2014;38: 89–98.

64. Stes A, Min-Leliveld M, Gijbels D, Van Petegem P. The impact of instructional development in higher education: The state-of-the-art of the research. Educational Research Review, 2009;5: 25–49.

65. Steinert Y, Mann K, Anderson B, Barnett BM, Centeno A, Naismith L, Prideaux D, Spencer J, Tullo E, Viggiano T, Ward H. A systematic review of faculty development initiatives designed to enhance teaching effectiveness: A 10-year update: BEME Guide No. 40. Medical Teacher, 2016;38: 142–159.

66. Chalmers D, Gardiner D. An evaluation framework for identifying the effectiveness and impact of academic teacher development programmes. Studies in Educational Evaluation, 2015;46: 81–91.

67. Austin AE. Faculty cultures, faculty values. New directions for institutional research. 1990;68: 61–74.

68. Trowler P, Bamber R. Compulsory higher education teacher training: Joined-up policies, institutional architectures and enhancement cultures. International Journal for Academic Development, 2005;10: 79–93.

69. Lund TJ, Stains M. The importance of context: an exploration of factors influencing the adoption of student-centered teaching among chemistry, biology, and physics faculty. International Journal of STEM Education, 2015;2: 13.

70. Austin AE. The socialization of future faculty in a changing context. The American academic profession: Transformation in contemporary higher education, 2011;145.

71. Pifer M and Baker V. Managing the process: The intradepartmental networks of early-career academics. Innovative Higher Education, 2013;38: 323–337.

72. Volkwein JF, Carbone DA. The impact of departmental research and teaching climates on undergraduate growth and satisfaction. The Journal of Higher Education, 1994;65: 147–167.

73. Martin E, Trigwell K, Prosser M, Ramsden P. Variation in the experience of leadership of teaching in higher education. Studies in Higher Education, 2003;28: 247–259.

74. Roxå T, Mårtensson K. Significant conversations and significant networks – Exploring the backstage of the teaching arena. Studies in Higher Education, 2009;34: 547–559.

75. Walter EM, Beach AL, Henderson CH, Williams CT. Describing instructional practice and climate: Two new instruments. In Weaver, G. C., Burgess, W. D., Childress, A. L., & Slakey, L. (Eds.), Transforming institutions: undergraduate STEM education for the 21st century. (pp. 411–428). Purdue University Press. 2015.

76. Landrum RE, Viskupic K, Shadle SE, Bullock D. Assessing the STEM landscape: the current instructional climate survey and the evidence-based instructional practices adoption scale. International journal of STEM education, 2017;4: 25.

77. Trigwell K, Prosser M. Development and use of the approaches to teaching inventory. Educational Psychology Review, 2004;16: 409–424.

78. Sawada D, Piburn MD, Judson E, Turley J, Falconer K, Benford R, Bloom I. Measuring Reform Practices in Science and Mathematics Classrooms: The Reformed Teaching Observation Protocol. School Science and Mathematics, 2002;102: 245–253.

79. Entwistle N, McCune V, Hounsell J. Approaches to study and perceptions of university teaching–learning environments: Concepts, measures and preliminary findings. Edinburgh, Scotland: Enhancing Teaching-Learning Environments in Undergraduate Courses Project, University of Edinburgh, Coventry University, and Durham University. 2002.

80. Kane R, Sandretto S, Heath C. Telling Half the Story: A Critical Review of Research on the Teaching Beliefs and Practices of University Academics. Review of Educational Research, 2002;72: 177–228.

81. Hemmings BC. Strengthening the teaching self-efficacy of early career academics. Issues in Educational Research, 2015;25: 1–17.

82. Bathgate ME, Aragón OR, Cavanagh AJ, Frederick J, Graham MJ. Supports: A key factor in faculty implementation of evidence-based teaching. CBE Life Sciences Education, 2019;18: 1–9.

83. Rosseel Y. Lavaan: An R package for structural equation modeling and more. Version0.5–12 (BETA). J. Stat. Softw. 2012;48, 1–36.

84. Harshman J, Stains M. A review and evaluation of the internal structure and consistency of the Approaches to Teaching Inventory. Int. J. Sci. Educ. 2017;39: 918–936.

85. Revelle W (2020). psych: Procedures for Psychological, Psychometric, and Personality Research. Northwestern University, Evanston, Illinois. R package version 2.0.7, 2020. https://CRAN.R-project.org/package=psych.

86. The Carnegie Classification of Institutions of Higher Education. About Carnegie Classification. Retrieved from http://carnegieclassifications.iu.edu/.

87. Barton K. MuMIn: Multi-model inference, R package (Version 0.12). 2009. From https://r-forge.r-project.org/projects/mumin

88. Burnham KP, Anderson DR. Model selection and multimodel inference: A practical information-theoretical approach. New York: Springer. 2002.

89. Grueber CE, Nakagawa S, Laws RJ, Jamieson IG. Multimodel inference in ecology and evolution: challenges and solutions. Journal of evolutionary biology, 2011;24: 699–711.

90. R Foundation for Statistical Computing, 161 Vienna, Austria, www.R-project.org/.

91. Chapman L, Ludlow L. Can downsizing college class sizes augment student outcomes? An investigation of the effects of class size on student learning. The Journal of General Education, 2010;2: 105–123.

92. Fung L, Chow LPY. Congruence of student teachers’ pedagogical images and actual classroom practices. Educational Research, 2002;44: 313–321.

93. Ebert-May D, Derting TL, Hodder J, Momsen JL, Long TM, Jardeleza SE. What We Say Is Not What We Do: Effective Evaluation of Faculty Professional Development Programs. BioScience, 2011;61: 550–558.

94. Woolfolk Hoy A. Self-efficacy in college teaching. Essays on teaching excellence: Toward the best in the academy, 2004;15: 8–11. Fort Collins, CO: POD Network. Retrieved March 23, 2020, from https://podnetwork.org/content/uploads/V15-N8-Woolfolk-Hoy.pdf

95. Ross JA. Teacher efficacy. In Hattie, J. & Anderman, E. M. (Eds.), International guide to student achievement (pp. 266–267). New York: Routledge. 2013.

96. Tschannen-Moran M, McMaster P. Sources of self-efficacy: Four professional development formats and their relationship to self-efficacy and implementation of a new teaching strategy. The Elementary School Journal, 2009;110: 228–245.

97. Major C, Dolly J. The importance of graduate program experiences to faculty self- efficacy for academic tasks. The Journal of Faculty Development, 2003;19: 89–100.

98. Hemmings B. Sources of research confidence for early career academics: A qualitative study. Higher Education Research & Development, 2012;31: 171–184.

99. Rienties B, Brouwer N, Lygo-Baker S. The effects of online professional development on higher education teachers’ beliefs and intentions towards learning facilitation and technology. Teaching and teacher education, 2013;29: 122–131.

100. Ebert-May D, Derting TL, Henkel TP, Maher JM, Momsen JL, Arnold B, Passmore HA. Breaking the Cycle: Future Faculty Begin Teaching with Learner-Centered Strategies after Professional Development. CBE-Life Sciences Education, 2015;14

101. Condon W, Iverson ER, Manduca CA, Rutz C, Willett G. Faculty development and student learning: Assessing the connections. Indiana University Press; 2016.

102. Cox MD. Faculty learning communities: Change agents for trans- forming institutions into learning organizations. To Improve the Academy, 2001;19: 69–93.

103. Owens MT, Trujillo G, Seidel SB, Harrison CD, Farrar KM, Benton HP., … Byrd DT. Collectively improving our teaching: attempting biology department–wide professional development in scientific teaching. CBE—Life Sciences Education, 2018;17: ar2.

104. Austin AE, Baldwin RG. Faculty Collaboration: Enhancing the Quality of Scholarship and Teaching. ASHE-ERIC Higher Education Report No. 7, 1991. ERIC Clearinghouse on Higher Education, George Washington University, One Dupont Circle, Suite 630, Washington, DC 20036.

105. Martin GA, Double JM. Developing higher education teaching skills through peer observation and collaborative reflection. Innovations in Education and Training International, 1998;35: 161–170.

106. Gehrke S, Kezar A. The Roles of STEM Faculty Communities of Practice in Institutional and Departmental Reform in Higher Education. American Educational Research Journal, 2017;54: 803–833.

107. Benbow RJ, Lee C. Teaching-focused social networks among college faculty: exploring conditions for the development of social capital. Higher Education, 2019;78: 67–89.

108. Withers M. The college science learning cycle: an instructional model for reformed teaching. CBE—Life Sciences Education, 2016;15: es12.

109. Hubball H, Collins J, Pratt D. Enhancing reflective teaching practices: Implications for faculty development programs. Canadian Journal of Higher Education, 2005;35: 57–81.

110. Green RG. Tenure and promotion decisions: The relative importance of teaching, scholarship, and service. Journal of social work education, 2008;44: 117–128.

111. Schimanski LA, Alperin JP. The evaluation of scholarship in academic promotion and tenure processes: Past, present, and future. F1000Research, 2018;7.

112. Gormally C, Evans M, Brickman P. Feedback about teaching in higher ed: Neglected opportunities to promote change. CBE—Life Sciences Education, 2014;13: 187–199.

113. Austin AE, Sorcinelli MD. The future of faculty development: Where are we going? New directions for teaching and learning. 2013;133: 85–97.

114. Kenyon KL, Cosentino BJ, Gottesman AJ, Onorato ME, Hoque J, Hoskins SG. From CREATE Workshop to Course Implementation: Examining Downstream Impacts on Teaching Practices and Student Learning at 4-Year Institutions. BioScience, 2019;69: 40–46.

115. Massy WF, Wilger AK, Colbeck C. Departmental cultures and teaching quality: Overcoming “hollowed” collegiality. Change: The Magazine of Higher Learning, 1994;26:11–20.

116. Feldman KA, Paulsen MB. Faculty motivation: The role of a supportive teaching culture. New directions for teaching and learning, 1999;78: 69–78.

117. Collie SL, Taylor AL. Improving teaching quality and the learning organisation. Tertiary Education & Management, 2004;10: 139–155.

118. Andrews TC, Conaway EP, Zhao J, Dolan EL. Colleagues as change agents: How department networks and opinion leaders influence teaching at a single research university. CBE—Life Sciences Education, 2016;15: ar15.

119. Thomson KE, Trigwell KR. The role of informal conversations in developing university teaching? Studies in Higher Education, 2018;43: 1536–1547.

120. Poole G, Iqbal I, Verwoord R. Small significant networks as birds of a feather. International Journal for Academic Development, 2019;24: 61–72.

121. Corbo JC, Reinholz DL, Dancy MH, Deetz S, Finkelstein N. Framework for transforming departmental culture to support educational innovation. Physical Review Physics Education Research, 2016;12: 010113.

