## Supplemental Materials for "Environmental influences and individual characteristics that affect learner-centered teaching practices"

**S1. Relevant Background Survey Questions**

*Please indicate the degree to which you agree or disagree with the following statements*

*(0=strongly disagree, 10=strongly agree).*

- My department is committed to transforming curricula and courses to enhance active learning and inquiry-based teaching.
- I frequently discuss issues pertaining to the improvement of teaching and learning with colleagues in my department.
- Other faculty in my department feel the same as I do about the need to improve undergraduate teaching and learning.
- Faculty in my department collaborate to achieve effective teaching (e.g., design, test, discuss curricula, etc.)
- Faculty in my department are interested in or are already conducting scholarly work about teaching and learning.
- Faculty in my department are recognized, evaluated, and rewarded for effective teaching.

*Please rate your knowledge of, first-hand experience with, and confidence about implementing each of the following.*

Knowledge (Low/Med/High), First-hand experience (Low/Med/High), Confidence (Low/Med/High)

- Course/curriculum planning
- Applying theories of learning (e.g., constructivism) to teaching practice
- Using technology in instruction
- Active learning
- Cooperative learning
- Using case studies
- Problem-based learning
- Inquiry-based teaching
- Assessment
- STEM education reform

*Imagine that you plan to develop and teach a course that uses active learning. Please use the scale below to indicate the degree to which the following would pose a challenge as you implement this course (1=not a challenge, 5=highly challenging). If an item is not applicable in your teaching context, please indicate that with the checkbox.*

1-5, N/A

- Time to plan, develop, and/or adapt materials
- Time to grade and/or give adequate feedback
- Time to train colleagues and/or TAs
- Cooperation/support of faculty in my department
- Cooperation of TAs and/or other instructors
- Support of campus administration
- Recognition or rewards for effective teaching
- Issues with reappointment/promotion
- Student attitudes toward and motivation around non-traditional teaching methods
- Student feedback through course evaluations
- Classroom infrastructure
- Access to instructional technology (e.g., clickers)

**S2. SCII items and exploratory factor analysis loadings for a six factor structure.**

Below are the items that loaded on each factor above 0.3. All loadings are in parentheses following each item.

Factor 1 - **Leadership**

- The department chair…
  - Encourages instructors to go beyond traditional approaches to teaching. (0.684)
  - Has a clear vision of how to improve teaching in the department. (0.826)
  - Implements teaching-related policies in a consistent and transparent manner. (0.767)
  - Inspires respect for his/her ability as a teacher. (0.807)
  - Is receptive to ideas about how to improve teaching in the department. (0.830)
  - Is tolerant of fluctuations in student evaluations when instructors are trying to improve their teaching. (0.588)
  - Is willing to seek creative solutions to budgetary constraints in order to maintain adequate support for teaching improvements. (0.757)

Factor 2 - **Mentoring and Material Support**

- Instructors in my department ARE…
  - Satisfied with their teaching workload. (0.444)
  - Assigned a mentor for advice about teaching. (0.322)
- Instructors in my department HAVE…
  - Adequate departmental funding to support teaching improvement. (0.656)
  - Adequate space to meet with students outside of class. (0.598)
  - Adequate time to reflect upon and make changes to their instruction. (0.739)
  - The support they need to employ educational technologies in their classrooms. (0.538)
- In my department…New instructors are provided with teaching development opportunities and resources. (0.386)

Factor 3 - **Flexibility in Teaching**

- Instructors in my department HAVE…
  - Considerable flexibility in the content they teach in their courses. (0.846)
  - Considerable flexibility in the way they teach their courses. (0.813)

Factor 4 - **Collegiality**

- Instructors in my department…
  - Frequently talk with one another. (0.909)
  - Discuss the challenges they face in the class-room with colleagues. (0.901)
  - Share resources (ideas, materials, sources, technology, etc.) about how to improve teaching with colleagues. (0.705)
  - Aspire to become better teachers. (0.403)

Factor 5 - **Evaluation of Effective Teaching**

- In my department…
  - Applicants for all teaching positions are required to provide evidence of effective teaching. (0.724)
  - Evidence of effective teaching is valued when making decisions about continued employment and/or promotion. (0.875)
  - Teaching effectiveness is evaluated fairly. (0.604)
  - Teaching is respected as an important aspect of academic work. (0.524)
  - All of the instructors are sufficiently competent to teach effectively. (0.482)

Factor 6 - **Support for Teaching Improvement**

- Instructors in my department…
  - Value teaching development services available on campus as a way to improve their teaching. (0.441)
- Instructors in my department ARE…
  - "Ahead of the curve" when it comes to implementing innovative teaching strategies. (0.430)
  - Willing to align the content of their courses to improve student learning. (0.361)
- In my department…There are structured groups organized around the support and pursuit of teaching improvement. (0.516)

**S3 Figure. Knowledge about teaching strategies.** These are faculty responding to their knowledge level with respect to active learning, assessment, and cooperative learning. The percentage is the percent of faculty reporting a certain level of knowledge within a group (FIRST IV or comparison).

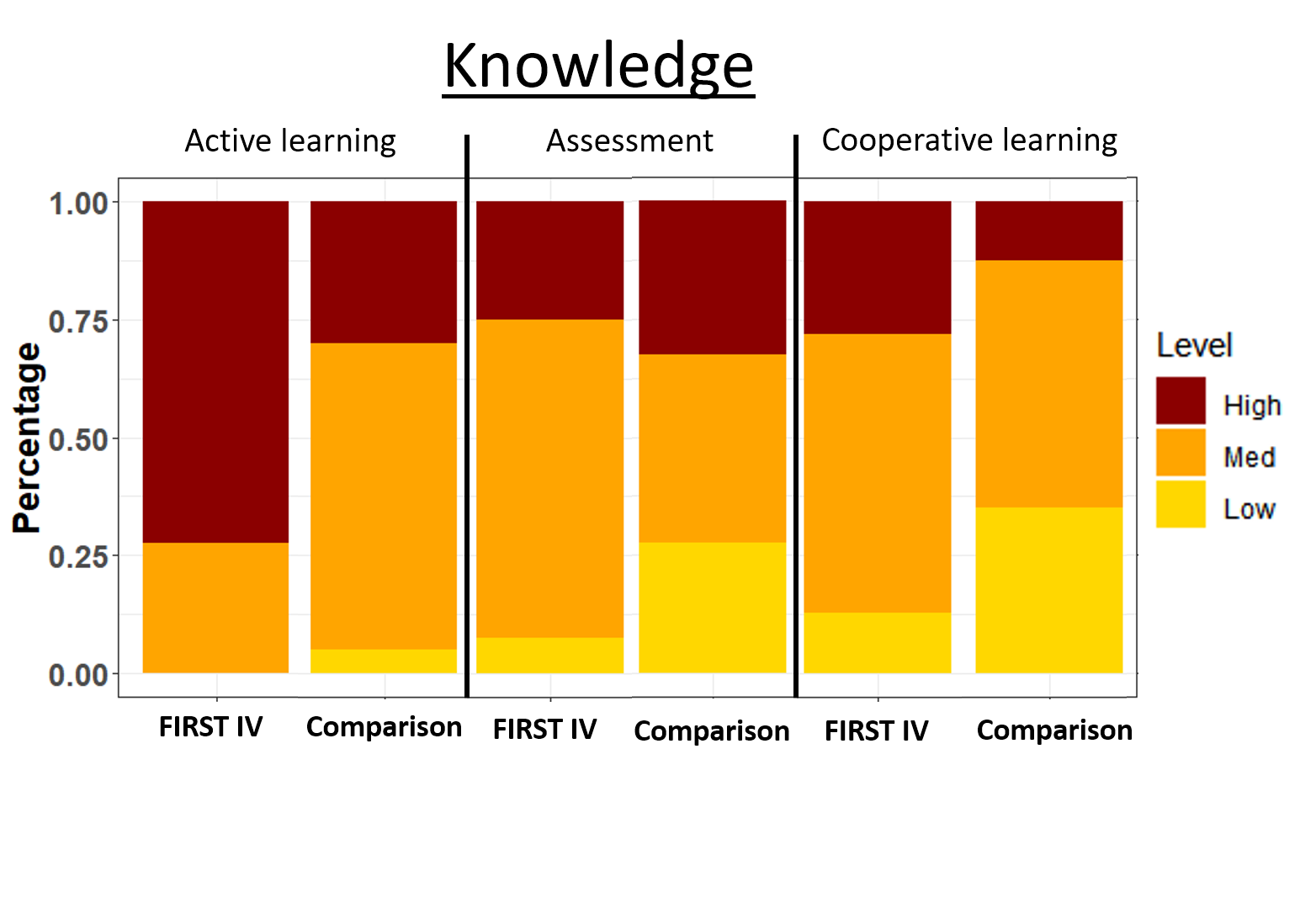

**S4 Figure. Experience about teaching strategies.** These are faculty responding to their experience level with respect to active learning, assessment, and cooperative learning. The percentage is the percent of faculty reporting a certain level of experience within a group (FIRST IV or comparison).

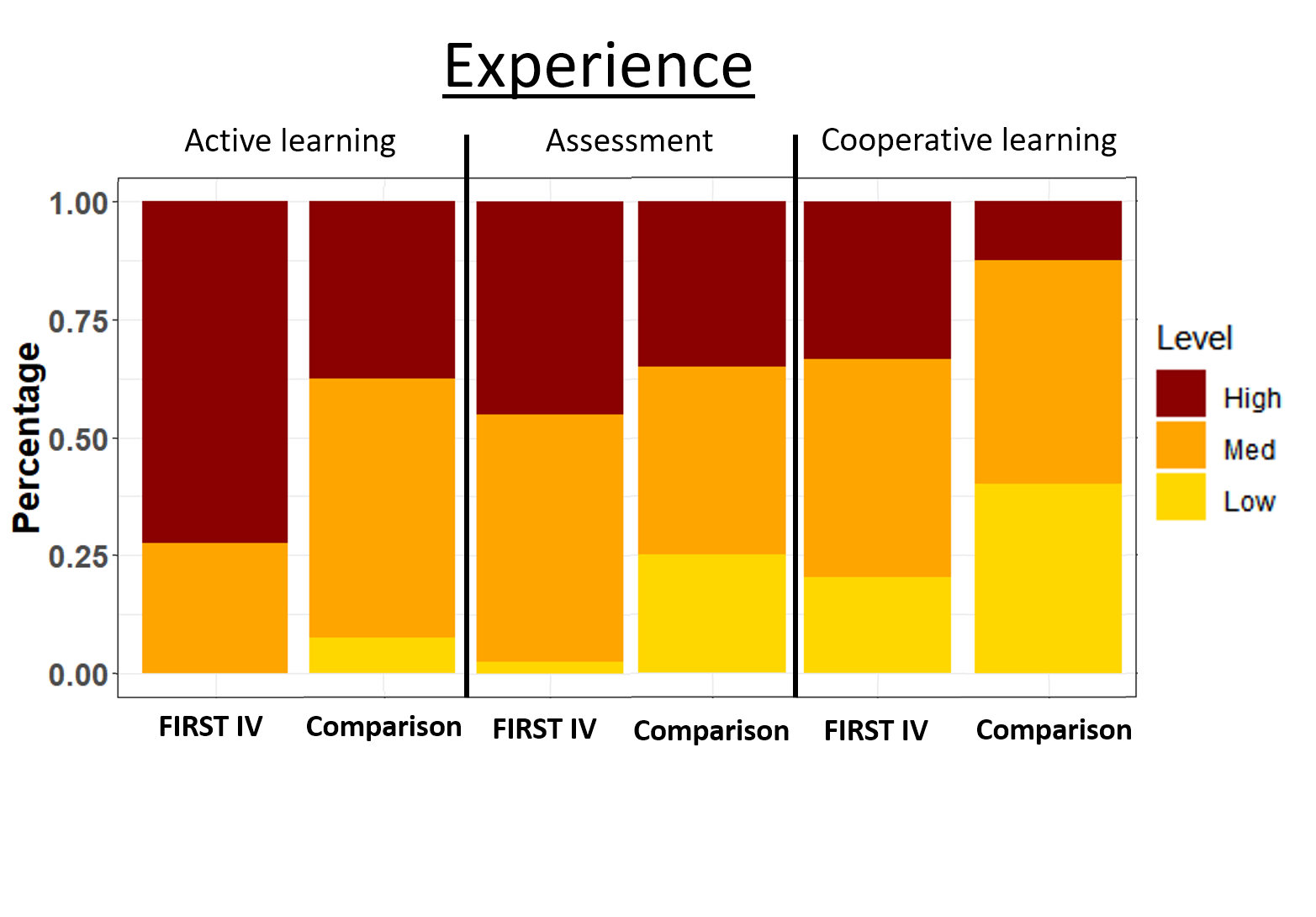

**S5 Tables. Model outputs.** AICc tables for DeltaRTOP & RTOP overall models

For Table 4

| **Change in RTOP Model Selection Table** | |  |  |  |  |  |  |  |  |
| --- | --- | --- | --- | --- | --- | --- | --- | --- | --- |
| **Intercept** | **Course Size** | **Percent Teaching** | **Self-Efficacy: Teaching Methods** | **Intentions: Knowledge Transmission** | **df** | **log-likelihood** | **AICc** | **delta** | **weight** |
| -0.014 |  |  | 0.332 |  | 3 | -41.885 | 90.7 | 0 | 0.285 |
| 0.13 |  |  |  |  | 2 | -43.514 | 91.5 | 0.8 | 0.192 |
| -0.085 |  |  | 0.337 | -0.243 | 4 | -41.011 | 91.6 | 0.9 | 0.182 |
| 0.063 |  |  |  | -0.237 | 3 | -42.77 | 92.4 | 1.77 | 0.116 |
| -0.027 | -0.126 |  | 0.362 |  | 4 | -41.479 | 92.5 | 1.84 | 0.114 |
| -0.033 |  | 0.162 | 0.332 |  | 4 | -41.513 | 92.6 | 1.9 | 0.11 |

For Table 5

| **Overall RTOP Model Selection Table** | |  |  |  |  |  |  |  |  |  |
| --- | --- | --- | --- | --- | --- | --- | --- | --- | --- | --- |
| **Intercept** | **Self-Efficacy:**  **Teaching Methods** | **Intentions:**  **Knowledge Transmission** | **Percent Teaching** | **FIRST IV / Comparison** | **SCII: Mentoring and Material Support** | **df** | **log-likelihood** | **AICc** | **delta** | **weight** |
| 0.2646 | 0.434 | -0.344 | -0.212 | + | 0.158 | 7 | -63.8 | 143.6 | 0 | 0.62 |
| 0.2561 | 0.454 | -0.361 | -0.189 | + |  | 6 | -65.53 | 144.5 | 0.98 | 0.38 |

**S6. Data Column Headers**

| **Column Header** | **Description** |
| --- | --- |
| IntID | Institution ID |
| PartID | Participant ID |
| PairID | Pair (FIRST IV & comparison) ID |
| FirstID | FIRST IV or comparison faculty |
| Course_YearID | Year of Study |
| Course_UID | Course unique ID |
| SCII_Factor1 | SCII Factor 1 - Leadership factor (derived from EFA) |
| SCII_Factor2 | SCII Factor 2 - Mentoring and Material Support |
| SCII_Factor3 | SCII Factor 3 - Flexibility in Teaching |
| SCII_Factor4 | SCII Factor 4 - Collegiality |
| SCII_Factor5 | SCII Factor 5 - Evaluation of Effective Teaching |
| SCII_Factor6 | SCII Factor 6 - Support for Teaching Improvement |
| Course_Size | Course enrollment |
| Percent_Research | Percent time allocated to research |
| Percent_Teaching | Percent time allocated to teaching |
| Agree_DeptTransform | Response to: Please indicate the degree to which you agree or disagree with the following statements (S1) |
| Agree_DiscussImprovement | Response to: Please indicate the degree to which you agree or disagree with the following statements (S1) |
| Agree_OtherFac_feel | Response to: Please indicate the degree to which you agree or disagree with the following statements (S1) |
| Agree_FacCollab | Response to: Please indicate the degree to which you agree or disagree with the following statements (S1) |
| Agree_FacInterest | Response to: Please indicate the degree to which you agree or disagree with the following statements (S1) |
| Agree_FacRewarded | Response to: Please indicate the degree to which you agree or disagree with the following statements (S1) |
| Knowledge_ActiveLearn | Knowledge level with respect to Active Learning |
| Knowledge_CoopLearn | Knowledge level with respect to Cooperative Learning |
| Knowledge_Assessment | Knowledge level with respect to Assessment |
| Experience_ActiveLearn | Experience level with respect to Active Learning |
| Experience_CoopLearn | Experience level with respect to Cooperative Learning |
| Experience_Assessment | Experience level with respect to Assessment |
| ALChallenge_TimePlan | Challenges to implementing active learning: Time to plan |
| ALChallenge_TimeGrade | Challenges to implementing active learning: Time to grade |
| ALChallenge_TimeTrain | Challenges to implementing active learning: Time to train colleagues/TAs |
| ALChallenge_FacCoop | Challenges to implementing active learning: Faculty cooperation |
| ALChallenge_TACoop | Challenges to implementing active learning: TA cooperation |
| ALChallenge_CampusSupport | Challenges to implementing active learning: Campus support |
| ALChallenge_TeachRewards | Challenges to implementing active learning: Teaching rewards |
| ALChallenge_Promotion | Challenges to implementing active learning: Promotion |
| ALChallenge_StudentAttitudes | Challenges to implementing active learning: Student attitudes |
| ALChallenge_StudentFeedback | Challenges to implementing active learning: Student feedback |
| ALChallenge_Room | Challenges to implementing active learning: Classroom infrastructure |
| ALChallenge_AccessTech | Challenges to implementing active learning: Access to technology |
| SE_CoursePlanning | Self-Efficacy subscale: Course planning |
| SE_TeachMethods | Self-Efficacy subscale: Teaching methods |
| SE_CreateLearnEnv | Self-Efficacy subscale: Creating a learning environment |
| SE_AssessStudentLearn | Self-Efficacy subscale: Assessing student learning |
| SE_InteractwStudents | Self-Efficacy subscale: Interacting with students |
| SE_MasterSubjKnow | Self-Efficacy subscale: Mastery of subject knowledge |
| TBI_B_LF | Teaching Beliefs & Intentions: Belief: Learning Facilitation |
| TBI_B_KT | Teaching Beliefs & Intentions: Belief: Knowledge Transmission |
| TBI_I_LF | Teaching Beliefs & Intentions: Intentions: Learning Facilitation |
| TBI_I_KT | Teaching Beliefs & Intentions: Intentions: Knowledge Transmission |
| RTOP_Score | Mean Reformed Teaching Observation Protocol score |
